# *In silico* modelling human VPS13 proteins associated with donor and target membranes suggests lipid transfer mechanisms

**DOI:** 10.1101/2022.07.27.501698

**Authors:** Filippo Dall’Armellina, Massimiliano Stagi, Laura E. Swan

## Abstract

The VPS13 protein family constitutes a novel class of bridge-like lipid transferases. Autosomal recessive inheritance of mutations in VPS13 genes is associated with the development of neurodegenerative diseases in humans. Bioinformatic approaches previously recognised the domain architecture of these proteins. In this study, we model the first ever full-length structures of the four human homologs VPS13A, VPS13B, VPS13C, and VPS13D in association with model membranes, to investigate their lipid transfer ability and potential structural association with membrane leaflets. We analyse the evolutionary conservation and physicochemical properties of these proteins, focusing on conserved C-terminal amphipathic helices that disturb organelle surfaces and that, adjoined, resemble a traditional Venetian gondola. The gondola domains share significant structural homology with lipid droplet surface-binding proteins. We introduce *in silico* protein-membrane models displaying the mode of association of VPS13A, VPS13B, VPS13C, and VPS13D to donor and target membranes, and present potential models of action for protein-mediated lipid transfer.

## Introduction

The ability of membranes to maintain organelle-specific physical properties, while responding to dynamic changes in cellular signalling and membrane flux requires a series of competing mechanisms of lipid synthesis, catabolism and homeostasis. Recent studies have shown that, apart from better-understood mechanisms of lipid synthesis, membrane sorting, trafficking and admixture to generate organelle-specific membranes, direct transfer of lipids from organelles, primarily the endoplasmic reticulum (ER), play a key role in cellular function and human disease.

Bidirectional interorganelle lipid exchange at mitochondria-associated ER membranes is regulated in yeast by the ER-mitochondria encounter structure (ERMES) complex (Jeong et al, 2016; Lang et al, 2015). Vacuolar protein sorting 13 (VPS13) is a family of universally conserved lipid transferase proteins that compensate for the absence of ERMES in metazoans (John Peter et al 2022, John Peter et al 2017). In yeast, *VPS13* is found as a single gene and was previously shown to localise at contact sites named v-CLAMPs (vacuole and mitochondria patch), and at nuclear-vacuole junctions known as NVJs (Lang et al., 2015). Recent studies predicted and analysed the full-length structural conformation of VPS13 in yeast (Toulmay et al, 2022), and hereafter, we focus on the architectural characterisation of human VPS13A, VPS13B, VPS13C, and VPS13D proteins in respect to their donor and target membranes.

The mechanisms of association between VPS13 proteins and organelle lipid membranes are as yet unclear. We undertake an *in silico* approach to investigate possible mechanisms of the VPS13 lipid transfer function. An N-terminal hydrophobic groove was previously found to solubilise tens of lipid fatty acid moieties (Adlakha et al, 2022; Kolakowski et al, 2021; Kumar et al, 2018) and extends across the bulk length of the protein in the form of β-strand repeats and their laterally connected α-helices or unstructured peptides, giving rise to the recently named repeating β-groove (RBG) domain (Neuman et al, 2022). Recognition motifs and protein-binding domains decorate VPS13s along their lipid transferring channels, which span across intermembrane contact sites that may vary between ∼10-40 nm in distance (Cai et al, 2022; Neuman et al, 2022). At the C-terminus, a VPS13 Adaptor Binding (VAB) domain interacts with a Pro-x-Pro motif to mediate protein interactions (Bean et al, 2018), in proximity to a pleckstrin homology (PH)-like domain (Fidler et al, 2016) which faces the cytosol. However, the VPS13 interactome is currently restricted to a few candidate physical partners mentioned below. The main structural characteristic of the VPS13s, is the presence of helical bundles at the C-terminal end where these lipid transferases associate with organelle membranes such as mitochondria and endo-lysosomes. These dual amphipathic helices (Kumar et al, 2018), we discover, are remarkably long and share common structural and biophysical properties with other lipid droplet surface-binding proteins, namely perilipins.

The significance of the loss or dysfunction of human VPS13 proteins is of increasing interest as this family is associated with neuropathogenic phenotypes. The mutational spectrum of VPS13A is associated with choreoacanthocytosis (Walker et al, 2007; Rampoldi et al, 2001). Choreoacanthocytosis is a rare autosomal recessive neurodegenerative disorder characterised by involuntary movements resembling Huntington’s disease, cognitive impairment, and abnormally shaped erythrocytes (Dobson-Stone et al, 2002; Dobson-Stone et al, 2005; Rampoldi et al, 2001). Interestingly, VPS13A is thought to act as an intermembrane bridge for the transfer of lipid cargo at ER-mitochondria, ER-lipid droplet, and mitochondria-endosome contact sites (Kumar et al, 2018; Li et al, 2020; Muñoz-Braceras et al, 2019; Ugur et al, 2020). Within its interactome, previous work found that VPS13A binds the lipid scramblase XK (Park and Neiman, 2020), Arf1 GTPase and *bis-* and *tris*-phosphorylated phosphoinositide lipids (Kolakowski et al, 2021). The closest paralog to VPS13A is VPS13C (Kumar et al, 2018), whose dysregulation, importantly, contributes to the progression and development of Parkinson’s disease (Lesage et al., 2016). VPS13C also localises at ER-lipid droplet, at ER-endo-lysosome jucntions (Kumar et al, 2018). Here, it is thought to interact with the GTPase Rab7 (McCray et al, 2010, Hancock-Cerutti et al 2022). Tethered at Golgi-endosome contact sites are VPS13B proteins (Seifert et al, 2011), which were shown to bind Rab6 (Seifert et al, 2015). The loss of functional VPS13B is pathogenic for Cohen syndrome pathology, a rare autosomal multisystem disorder principally characterised by physiological defects, a distorted perception of self, intellectual disability, and intermittent congenital neutropenia (Duplomb et al, 2014; Kolehmainen et al., 2003). Finally, a newly described form of spastic ataxia was associated with mutations in *VPS13D* (Gauthier et al., 2018; Seong et al., 2018). The largest protein in the family, VPS13D is localised at peroxisomes and Golgi complex (Baldin et al, 2021; Guillén-Samander et al, 2021). Recently, it was found that the GTPases Miro1 and Miro2, orthologues of the outer mitochondrial membrane GTPase Gem1 which interacts with ERMES in yeast, also recruit VPS13D at contact sites between mitochondria and the ER (Guillén-Samander et al, 2021).

To understand better how human VPS13 proteins function during exchange of lipids between organelles, and to explore their dysfunction in disease, we made an in sillico analysis of the protein architecture, evolutionary conservation and biophysical properties unique to each human paralog. We note several domain features which are unique to each VPS13 protein, along with islands of conservation indicative or protein-protein interaction partners. We find that all VPS13 proteins share lipid docking properties similar to perlipins, but that for each of the VPS13 paraologs, a perilipin-C-like domain, which we name the gondola domain, associates with target membrane outer leaflets, with paralog-specific properties, to facilitate lipid transfer. We thus present two possible models of passive lipid transfer *via* VPS13 family proteins.

## Methods

### Protein modelling

The transform-restrained Rosetta server was employed to predict the structural conformation of our proteins of interest (Du et al, 2021; Su et al, 2021; Yang et al, 2020). Prior to any webserver submissions, we ran multiple sequence alignments for the canonical transcripts of VPS13A (NP_150648.2), VPS13B (NP_060360.3), VPS13C (NP_065872.1), and VPS13D (NP_056193.2). These were executed meeting the following requirements, Python 3.7.13, Tensorflow 1.14.0, PyRosetta4, Perl 5.32.1, HH-suite3.3.0, Uniclust compilation (available at http://www.user.gwdg.de/~compbiol/uniclust/2022_02/), and the trRosettaX package. The number of alignments used for modelling was capped at 1,500 hits closest to the query. Separate FASTA files containing regions of 1,500 amino acids were created to cover our full-length proteins. Furthermore, sequence-dependent alignment and structural superimposition of the separate models for each individual protein were performed on PyMOL (Schrödinger and DeLano, 2020), where each selection contained ∼420 amino acid residues with 100% identity to its precedent and subsequent regions. For VPS13B, we used the longest transcript, however, we note that studies that localised VPS13B the Golgi complex examined the splices isoforms NM_152564.3, NP_689777.3.

### Evolutionary conservation analysis

The VPS13 family proteins were searched on the NCBI Orthologs Database and one protein sequence per gene was aligned with a constraint E-value of 0.003 on COBALT (available at https://www.ncbi.nlm.nih.gov/tools/cobalt/re_cobalt.cgi), allowing the programme to identify conserved columns and recompute the alignment. The multiple sequence alignments generated for VPS13A, VPS13C, and VPS13D, were taken from subphylum *Gnathostomata* (jawed vertebrates) and the VPS13B sequences were taken from *Craniata* (vertebrates), where each alignment group comprised around 300 species. Subsequently, the ConSurf Server was employed to display the conservation colour-coding scheme of our proteins (Ashkenazy et al, 2016; Landau et al, 2005).

### Surface hydrophobicity

The Eisenberg scale (available at https://web.expasy.org/protscale/pscale/Hphob.Eisenberg.html) was employed to investigate the hydrophobicity of the structure predictions created with trRosetta. This amino acid scale is a normalized consensus hydrophobicity scale that assigns specific values to each residue. The color_h.py PyMOL command was finally run to colour-code our single chain models.

### Determination of electrostatic potentials

The electrostatic potentials of each protein were calculated using the APBS Electrostatics Plugin in Incentive PyMOL 2.0 on the APBS-3.4.1 version of the software (Jurrus et al, 2018) and were displayed as colour-coded surfaces. The PDB2PQR programme was implemented as default for processing partial charges and atom radii where calculations were performed at 0.15 M ionic strength (ion charge +1 with solvent radius 2, ion charge −1 with solvent radius 1.8, 310 Kelvin, pH 7, protein dielectric 2, and solvent dielectric 78). The PARSE forcefield was run on the prepared models and the electrostatic potential surfaces were displayed on a red-white-blue colour map with a range of +/- 6.00 kJ/mol/e.

### Structural pairwise alignments

The C-termini of our VPS13s models were submitted to BackPhyre to scan the Homo sapiens genome for similar structures (Kelley et al, 2015). The structure comparison platform FATCAT (Flexible structure AlignmenT by Chaining Aligned fragment pairs allowing Twists) was implemented to predict the conformational similarity of our model chains (Ye and Godzik, 2003) against TIAM1 (T-cell lymphoma invasion and metastasis-inducing protein 1, AlphaFold AF-Q13009-F1-model_v2; UniProt Q13009), ATG2A (Autophagy-related protein 2 homolog A, AlphaFold AF-Q2TAZ0-F1-model_v2; UniProt Q96BY7), Plin2 (Perilipin 2, AlphaFold AF-Q99541-F1-model_v2, UniProt Q99541), and GPAT4 (Glycerol-3-phosphate acyltransferase 4, AlphaFold AF-Q86UL3-F1-model_v2; UniProt Q86UL3).

### Simulation of lipid membrane-protein interactions

Protein-lipid membrane simulations were executed using the CHARMM-GUI membrane builder (Brooks et al, 2009; Jo et al, 2008, Wu et al, 2014) with CHARMM36 lipid force field (Klauda et al, 2010). PPM 2.0 was run on the input protein models to orientate the molecular structures submitted in reference to the lipid leaflets. The force field options assigned were CHARMM36m/WYF parameter for cation-pi interactions/HMR and NAMD for the input generation. The system equilibration and relaxation used the automatic PME FFT grid generation method and NPT ensemble for lipid bilayers at a temperature of 310.15 K (Wu et al, 2014). Finally, protein-lipid membrane components were assembled through the CHARMM-GUI Replacement Method. Lipid membranes compositions were based on previous predictions and biochemistry (Escribá et al, 2015; Guo et al, 2009; Horvath and Daum, 2013; Jo et al, 2008; Pogozheva et al, 2022; Reglinski et al, 2020; Tauchi-Sato et al, 2002; van Meer and de Kroon, 2011; van Meer et al, 2008). Protein-membrane orientations were based on previous experimental approaches (Baldwin et al, 2021; Koike et al, 2019; Kumar et al, 2018; Muñoz-Braceras et al, 2019; Seifert et al, 2011; Yeshaw et al, 2019).

## Results

### Structural Architecture of VPS13s

The domain architecture nomenclature of each protein was adapted from previous literature (Fidler et al, 2016; Kumar et al, 2018; Leonzino et al, 2021). As expected, our modelled human VPS13 formed a Chorein domain at the N-terminus, followed by an extended region of β-sheets with a highly hydrophobic interior, known as the repeating β-groove (RBG) domain (Neuman et al, 2022), which makes the bulk of the length of the protein (**Video 1**, **Fig1**). Our modelling shows that VPS13A, VPS13C, and VPS13D adopted an open conformation of the RBG domain where the hydrophobic surface interior, assumed to transport lipid cargoes, is partially exposed to solvent through a seam (Fig. 3). Conversely, the predicted structure of VPS13B adopted a more compressed conformation, somewhat reducing solvent exposure along the RBG channel, and was consequentially overall 20 Å shorter (**Video 1**).

**Figure 1.**
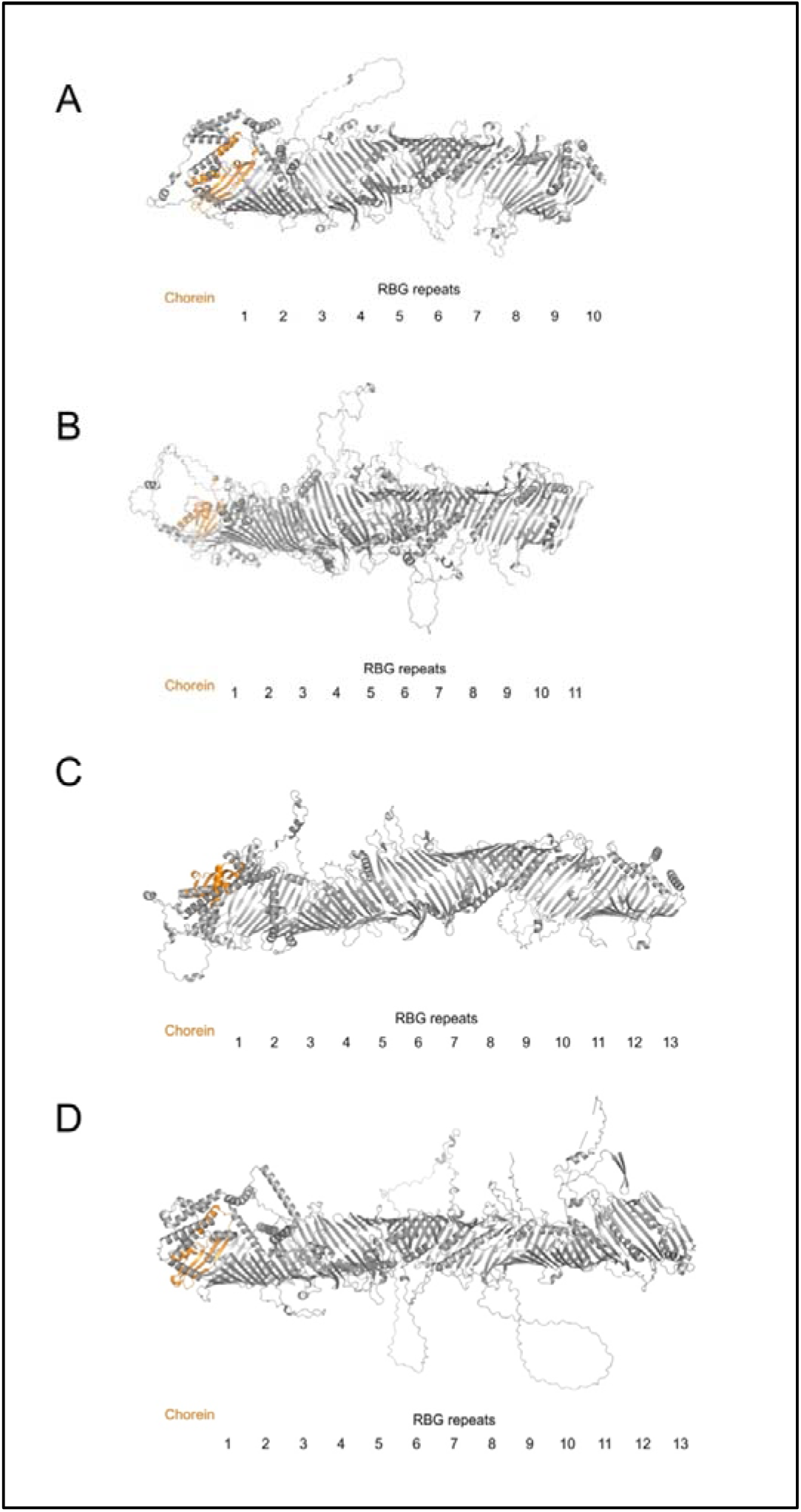
Architecture of the RBG domains in human VPS13 proteins (A-D). Chorein domain (orange), RBG repeats (grey). The models were studied in PyMOL (Schrödinger and DeLano, 2020).

At the N-terminal region, three of the four human VPS13 proteins display a two phenylalanines in a solvent-exposed acidic tract (FFAT) motif, which binds VAMP-associated proteins (VAP) tethering the ER (Murphy and Levine, 2016). The FFAT motif is located on a loop in VPS13A and on a short α-helix of the outer surface of the RBG repeats in VPS13C and VPS13D (**Video 1**). However, this interactor motif is lacking in VPS13B (NP_060360.3) and the region where we would expect it to be located on the protein is not highly conserved (**Video 2**). Uniquely, VPS13D (NP_056193.2) is the only homologue that expresses a ∼43 residue-long ubiquitin-associated (UBA) domain. The VPS13D UBA domain is a cytosolic-facing structure that comprises three short α-helices, precedes a loop and is located ∼50 nm away from the RBG outer sheets, potentially to allow space for an interactor protein to bind (**Video 1**).

It is currently believed that lipids transferred by VPS13 family protein are transported from a donor membrane, which abuts the N-terminal Chorein domain, to a target membrane defined by interactions of their C-terminal domains. Several proteins of the presumptive target membranes are known to interact with the C-terminal VPS13 Adaptor Binding (VAB) domain, previously known as SHR-BD (**Video 1**; Bean et al, 2018). Approximately 100 amino acid residues long, the VAB domain repeats share common structural characteristics to the bladed sheets of β-propeller and WD40 domains (Kumar et al, 2018). However, the VAB domain of VPS13 proteins lack the characteristic “WD” dipeptide repeats of WD40 domains and do not assemble as a donut-like toroid with a central pore (Mittal et al, 2018; Sprague et al, 2000) (**Fig.2**). On the other hand, the VAB domain contains six β-bladed repeats characterised by the presence of a highly conserved asparagine residue (**Video 2**; Bean et al, 2018). This residue is present in the β-sheet repeats of the VAB domains in all human VPS13 proteins, however, it is substituted for a serine in the second and third VAB domain repeats of VPS13B (**Video 1**; Bean et al, 2018; Dziurdzik et al, 2020; Dziurdzik and Conibear, 2021). In VPS13s, the VAB β-repeats are wrapped obliquely around the tube formed by the RBG repeats, and do not touch the inner surface of those RBG repeats, suggesting that the VAB domain serves to either regulate or recruit VPS13s to their target membrane *via* protein partners, rather than to directly interact with transferring lipids (**Video 1**).

**Figure 2.**
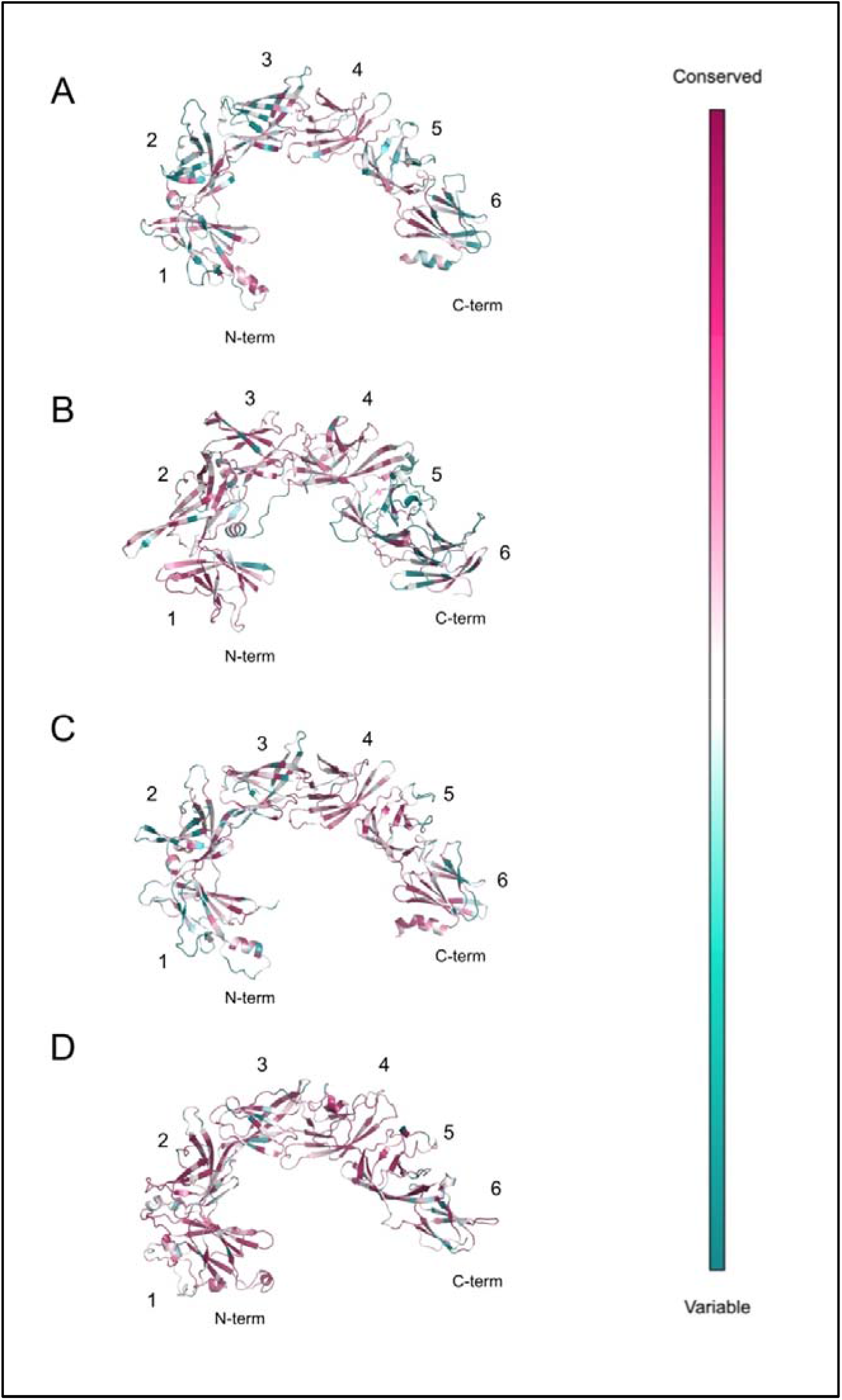
Evolutionary conservation of the VAB domains in human VPS13s (A-D). The six β-bladed repeats were labelled accordingly (Bean et al, 2018). The analysis was carried out employing the ConSurf Server (Ashkenazy et al, 2016; Landau et al, 2005) and the models were viewed in PyMOL (Schrödinger and DeLano, 2020). Darker purple-coloured residues are highly conserved and blue residues are variable in evolution.

Over the four orthologues, the general structure of the protein is largely similar as described above, however at the C-terminal portion of the proteins, several features were identified, which were unique to each orthologue. Our modelling revealed all four proteins contain a previously undescribed globular domain of elusive function that was found at one or other end of the VAB domain (**Video 1**), which we call for convenience the small globular domain (SGD). The SGD in VPS13s is a subregion of what has previously been named the APT1 domain (PF10351) at the C-terminus of yeast VPS13 and human VPS13A proteins (Dziurdzik and Conibear, 2021), which we will discuss in more detail below.

The SGD was divergent between each human VPS13 protein, but the closest and most significant structural homologs belonged to the polypeptide N-acetylgalactosaminyltransferase (pp-GalNAc-T) family (**Tables S1-S4**; Kelley et al, 2015). Rather than adopting a similar conformation to catalytic subdomains A and B of this protein family, the SGD of VPS13D fits the structural fold of their characteristic ricin B-type lectin domain, which is thought to bind to GalNAc and contribute to the glycopeptide specificity of pp-GalNAc-T proteins, potentially indicating VPS13 proteins associate with carbohydrate-containing proteins or participate in transport ofcarbohydrates (Velayos-Baeza et al, 2004). While the SGD fold of VPS13A is not particularly similar to a ricin B-type lectin domain, it is structurally homologous to WWE domains that are often found in ubiquitin ligases and may direct protein interactions (**Tables S1-S4**; Aravind, 2001).

Each of the four proteins display a C-terminal pleckstrin homology (PH)-like domain generally characterised by an α-helix and anti-parallel β-sheets (**Video 1**; Fidler et al, 2016). In our modelled PH-domains, several neighbouring helices and peptide loops are strongly conserved over evolution, and we included them in our definition of the VPS13 PH-like domain due to the significance of their evolutionary conservation (**Video 2**). The loops between the β-sheets of these PH-like domains differ in length in the different VPS13 proteins, but the anti-parallel topological orientation of the β-sheets is consistent across all human VPS13 proteins.

Most strikingly, the α-helical domain of the VPS13 C-terminus, described as a DH-like sequence or split into a region covered by the APT1 and ATG-C terminal homology domains (Dziurdzik and Conibear, 2021; Kumar et al, 2018), assumes a remarkable conformation, suggesting a key function in the lipid transfer action of VPS13 family proteins and structural paralogs such as ATG2 (**Tables S5-S8**, Velikkakath et al, 2012; Tamura et al, 2017). In all four proteins, this α-helical region extends as a bundle of giant helices that project beyond the plane formed by the VAB domain arch, the base of the tube formed by the RBG domain, and the hydrophobic β-sheets of the putative APT1 domain (Video 1; Fig. 3; Kumar et al, 2018).

**Figure 3.**
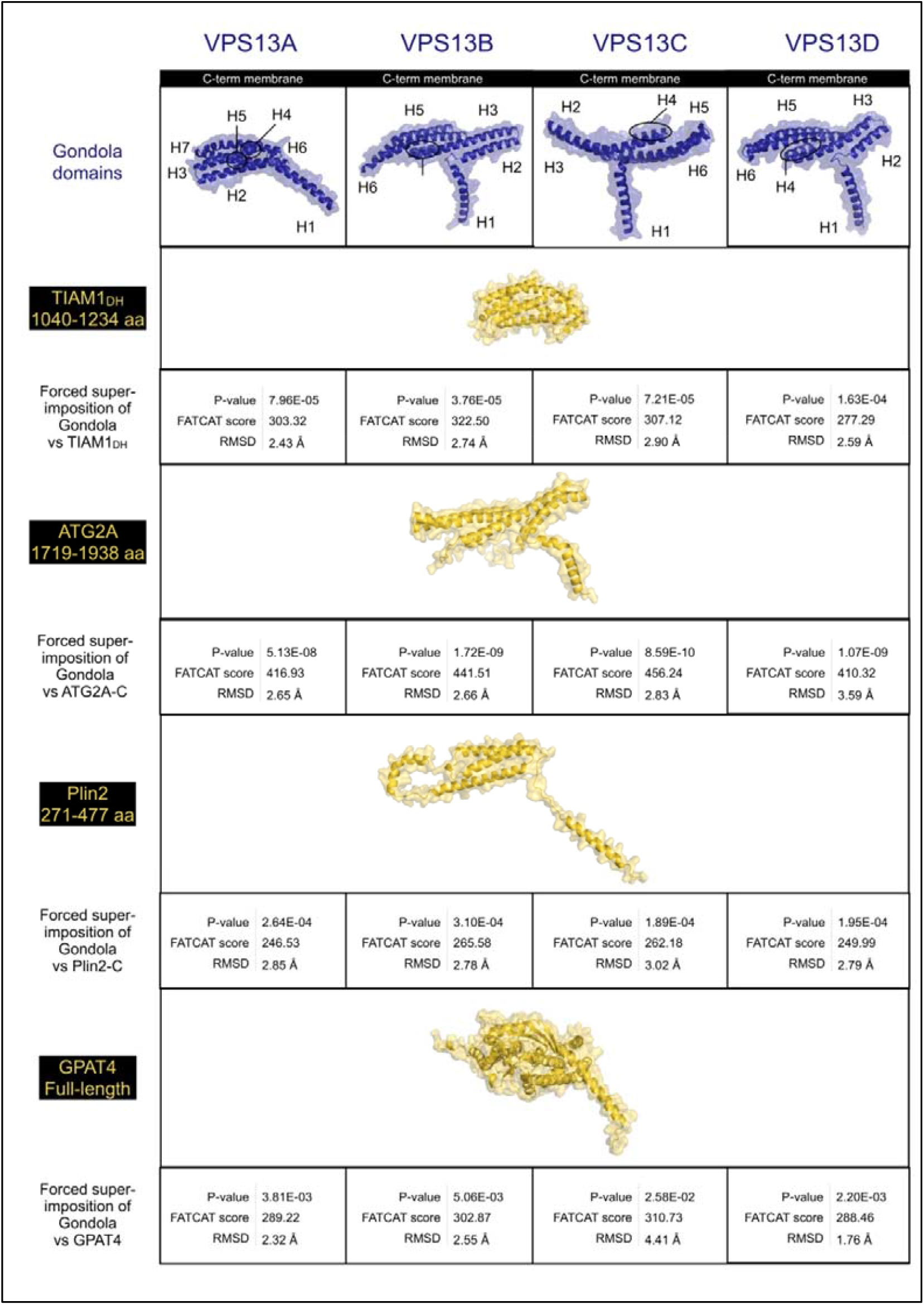
Structural homology of the gondola domains of human VPS13 proteins (A-D) with lipid-droplet coat proteins. VPS13 gondola (dark blue), other proteins (gold). Similarity scores obtained *via* FATCAT (Ye and Godzik, 2003).

As previously noted, these helices are amphipathic (Kumar et al, 2018; Rowe et al, 2016) and in our simulations of interaction with target membranes (see below), they directly insert into the outer leaflet of the target membrane, disordering the lipid surface (**Fig. 4**). Such giant dual amphipathic helical bundles remarkably resemble a traditional Venetian gondola and in fact we propose to name these structures the protein’s “gondola” domain, thus further subdividing the previous APT1-ATG-C homology region. The rationale behind this is that, despite the sequence and structural homology of the typical APT1 domain (PF10351) with the SGD-gondola region of VPS13s, the VPS13 APT1-like domains are characterised by short β-strands and α-helices that fold into a globular domain (part of SGD), more than five β-sheets similarly oriented to the ones of the RBG repeats (part of RBG), relatively few α-helices that similarly to the RBG domain connect laterally these β-strands (part of RBG), and a giant helix perpendicular to the inner hydrophobic lipid transporting pore of the protein that belongs to the gondola domain, as like the other helices of the gondola domain, it is of both extended length and amphipathicity.

**Figure 4.**
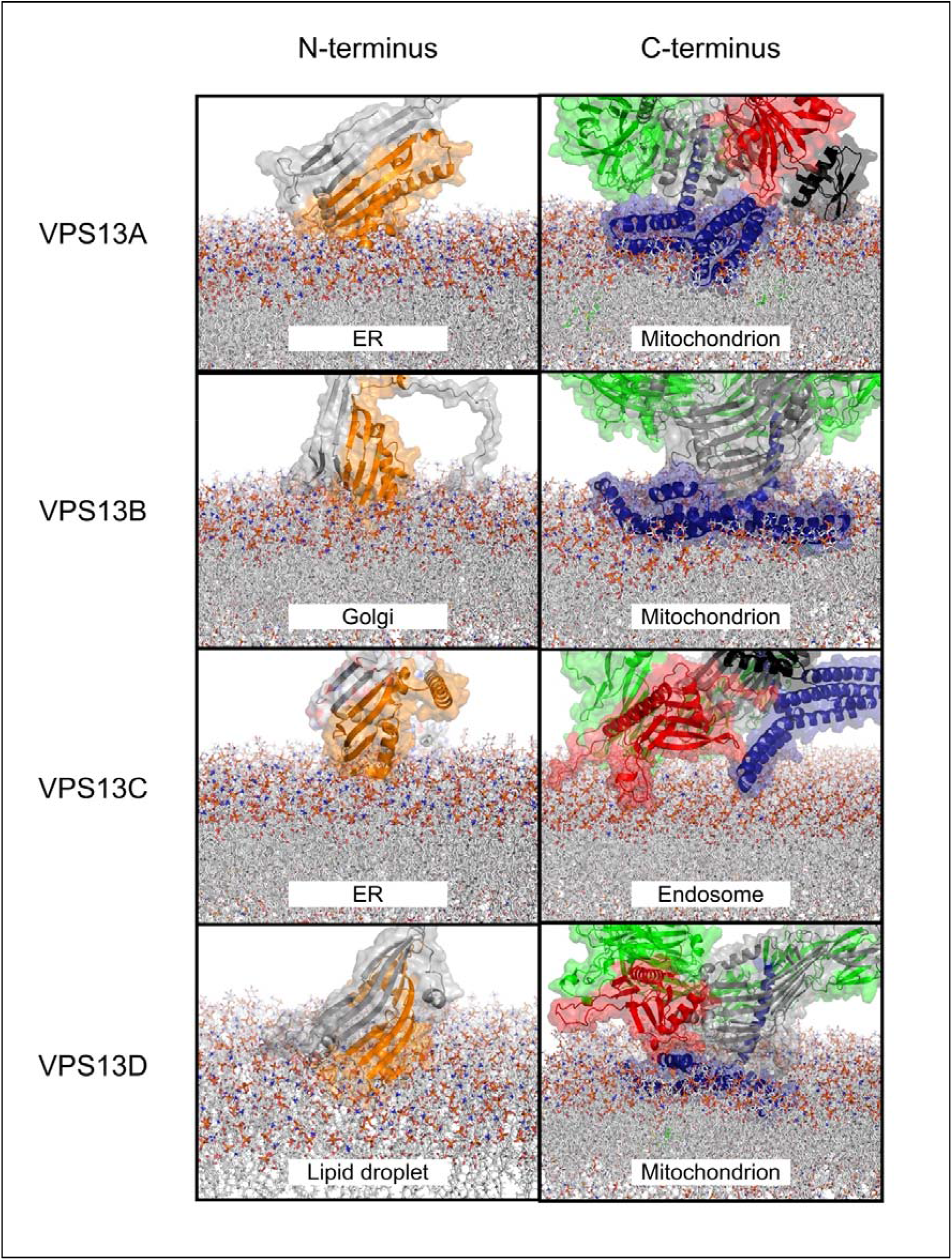
Association of human VPS13 proteins (A-D) with simulated donor and target membranes. *In silico* predictions were executed via CHARMM (Brooks et al, 2009; Jo et al, 2008, Wu et al, 2014). Chorein domain (orange), FFAT motif (magenta), RBG domain (grey), UBA domain (azure), VAB domain (green), SGD (black), gondola domain (dark blue), PH-like domain (red). Lipid leaflet compositions are reported in Tables S10-S18.

Importantly, the conformation of the VPS13A, VPS13B, VPS13C, and VPS13D gondolas significantly resembled structures of proteins bound at lipid droplet surfaces (P-values < 0.05 **Fig. 3; Tables S5-S8**). Pairwise alignments of our full-atom models discovered that human VPS13 gondola domains contain helices with similar lengths and angles to the ones seen in ATG2A, Plin2, and GPAT4 (**Fig. 3**). Previously, sequence homology has suggested that this domain may be a DH-like domain (Kumar et al, 2018), so we also modelled the C-terminal human VPS13 gondola domains against a typical DH domain (**Fig. 3**) taken from TIAM1 (UniProt Q13009). While there was some similarity, several points of divergence arose with TIAM1_DH_ such as that the VPS13 gondola domains are not characterised by the typical 11-helix bundle of DH domains but rather contain 6-7 giant helices that adopt a curved conformation (**Fig. 3**), as do the lipid binding domains of ATG2A, Plin2 and GPAT4. The N-terminal α-helix (H1) of the VPS13 gondola domain is perpendicular to the other helices, similarly to the bundle conformations seen in Plin2, and it is in fact oriented at an oblique angle to the target membrane (**Figs. 3, 4**). In our models, the gondola domain’s first α-helix (H1) points directly toward the interior of the RBG domain, partially occluding the end of the VPS13 β-groove hydrophobic inner channel, whose C-terminus is located just underneath the VAB domain, and bringing the pore size for lipids to travel through down to 1-2 nm, perhaps limiting or directing flow of lipid.

### Evolutionary Conservation of VPS13s

Conservation analysis was based on the sequence similarity of each human VPS13 within over 300 orthologues from vertebrate lineages. Significant conservation was seen at the N-terminal Chorein and RBG domains of VPS13B, and at the C-terminal gondola domains of all human VPS13s (**Video 2**). Along the length of the RBG domain of each VPS13 protein, several of the unstructured loops displayed well-conserved residues, as well as cytosolic-facing surfaces of the VAB and PH domains, that appear to be opposed to one another (**Video 2**, **Figure 2**), initially suggesting protein interaction partners along the length of VPS13s. These short peptide regions did not display any obvious interaction motifs and thus we did not further analyse them. The VAB domains appeared evolutionary variable regions, except for the N-terminal region of the VAB domain in VPS13B and for most cytosolic surface-facing residues in the VPS13D VAB domain (**Video 2, Fig. 2**). In VPS13A, the SGD was not conserved in terms of amino acid identity, whereas in the β-sheets of SGD in VPS13B and VPS13D, and in the SGD residues facing the RBG domain in VPS13C, the conservation was significant (**Video 2**).

We then mapped known missense mutations of the VPS13 family proteins (**Table S9**). Mutations were found in most domains across the VPS13 proteins. Importantly, three missense mutations in VPS13D sit at helical membrane-bound regions of the gondola domain and are in highly conserved residues, potentially suggesting their ability to disrupt protein docking of the C-terminus at the lipid membrane. Two of these mutations were deemed likely pathogenic (ClinVar VCV000932894.1, VCV000983263.1), and seem to cause autosomal recessive cerebellar ataxia-saccadic intrusion syndrome, whereas the third mutation (p.Ala4210Val, rs746736545, VCV000561200.1) causes spastic ataxia (Seong et al, 2018). Furthermore, multiple inward-facing missense mutations were found within the hydrophobic interior of the RBG domain, one of which was pathogenic for Cohen syndrome in VPS13B (p.Ile1636Asn, VCV000977843.1), and another one was likely pathogenic in VPS13D (p.Ile1170Ser, VCV000806071.11). It is possible that these mutations hinder lipid cargo transfer regulation in their respective disorders.

### Biophysical Properties of VPS13s

It was expected that the inner channel of the RBG domain in VPS13 proteins would be highly hydrophobic throughout (**Video 3**; Kumar et al, 2018; Leonzino et al, 2021). Suggestive of their association with donor and target membranes, both the N-terminal extremity of the Chorein domain and the C-terminal gondola domain are strongly hydrophobic. Interestingly, the orientation of the hydrophobic surface of the gondola domain is distinct between the four proteins. As previously described (Kumar et al, 2018), the gondola domain is composed of a series of amphipathic helices. In VPS13A, the most hydrophobic residues of the gondola domain are located between its two major α-helix bundles (**Video 3**). In VPS13B and VPS13D, the wing that is predicted to bind the membrane is highly hydrophobic, whereas in VPS13C, the amphipathicity of the gondola domain is inverted, such that the hydrophilic surface of the gondola appears oriented towards the target membrane and the hydrophobic surface orients toward the C-terminal tube of the RBG domain (**Video 3**).

We also simulated the charge distribution over the proteins. The gondola domain displays oppositely charged wings (one positive, one negatively charged), as seen in VPS13A and VPS13C (**Video 4**). However, these C-terminal regions appeared relatively neutral in VPS13B and VPS13D (**Video 4**). Potential interactor-docking pockets close to the plane of the membrane-bound gondola domain were also positively charged and in proximity to the RBG pore (**Video 4**).

### Protein-membrane dynamics

Our protein-membrane interaction simulations revealed the positioning of the human VPS13s bridging their presumptive donor and target membranes (**Fig. 4**). We considered the insertion of the protein into the membrane successful exclusively when residues resided below the phospholipid plane, and disturbed the outer leaflet of the modelled target membrane. In our modelling, the N-terminal β-sheets of the Chorein domain and the C-terminal gondola domain inserted below the plane of the lipid membrane surfaces (**Fig. 4, Videos S1-S5**). While the N-terminal Chorein domain associates shallowly with the ‘donor’ membrane outer leaflet, such that the hydrophobic ‘scoop’ is aligned perpendicularly to the membrane surface, the gondola domains are much more deeply pressed into the target membrane outer leaflet (**Fig. 4**). The VAB and PH-like domains and their conserved patches, which were not particularly electrostatically charged nor hydrophobic, were instead placed proximate to the target membrane surface without inserting into the membrane (**Fig. 4, Videos 3-4**). Interestingly, the only exception to this was the model of VPS13C at the endosomal membrane (**Video S3**) where, due to the inversion of the amphipathic surfaces of the VPS13C gondola (**Videos 3-4**), only one end of the gondola penetrated the membrane (**Fig. 4**). However, in this model, the loops that connect laterally the β-sheets of the VPS13C PH-like domain appeared to partially displace the lipid surface (**Fig. 4**) and in fact contained moderately hydrophobic residues (**Video 3**). This mechanism may assist in docking VPS13C to its target. Notably, in our models where the gondola domains penetrated the outer membrane leaflet of our target organelles, the pore of the tube formed by the RBG domain becomes closely opposed to the organelle surface, proximate to the region of the membrane that is disorganised due to the insertion of the gondola domain (**Videos S1-S4**).

## Discussion

### Insights in the VPS13 Chorein and RBG domains

How VPS13 proteins are able to transfer lipids and how they may be regulated, are currently unknown, despite significant recent interest in their function. Starting at the N-terminus, the four human VPS13s display a Chorein domain, which is also found in their yeast orthologue and ATG2 proteins (Chowdhury et al, 2018; Kumar et al, 2018; Lees and Reinisch, 2020; Valverde et al, 2019). The Chorein domain spans ∼120 amino acids and was recently discovered to be closely related to the N-termini of bacterial proteins AsmA and TamB and fungal Mdm31p, which are all predicted to contribute to lipid transfer (Levine, 2019). Two main features of the VPS13 Chorein domain were characterised through the crystallisation of a yeast VPS13 N-terminal 150 kDa fragment (PDB ID: 6CBC). In our modelling of ER and lipid droplet membranes, an open part of the extreme N-terminus dips into the membrane (**Fig. 4**), suggesting that there is not a great deal of selectivity for cargo at the N-terminal end of these proteins. The Chorein domain is formed by this scoop and one or two α-helices (Kumar et al, 2018), which are predicted to be present in a number of putative lipid transfer proteins such as ATG2, SHIP164, Csf1, and the Hob proteins (Neuman et al, 2022).

Over the length of the RBG domain, our conservation and structural models yielded the following observations. Namely, there were a surprising number of short strongly conserved peptides in unstructured loops which decorated the length of the RBG tube (**Fig. 1, Video 2**). In addition to another group of structured domains that we collectively called SGDs (**Video 1**), which appear closely related between VPS13A and VPS13C (Kumar et al, 2018), we found several conserved helical structures at the edges of the seam of the VPS13 proteins, potentially suggesting that these are embedded in a network of interactors which extends over the intramembrane bridge. Most curiously, such an architectural geometry characterised by RBG repeats and their laterally connected cytosolic loops appears to be a common feature of the RBG protein superfamily, especially in ATG2, Hob and SHIP164 (Hanna et al, 2022; Neuman et al, 2022).

We found no obvious peptide motifs in the conserved loops that decorate the length of the RBG domain. However, we discuss other accessory domains below and their possible role in regulating VPS13 proteins. We located a well conserved FFAT motif between residues 476-482 in VPS13D (**FFD**PTA**D**), which was cytosolic-facing in an alpha-helix surrounded by unstructured regions (**Video 1**; Leonzino et al, 2021; Di Mattia et al, 2020). This motif, generally consisting of around six amino acids, interacts with the major sperm protein (MSP) domain of vesicle-associated membrane protein (VAMP)-associated proteins (VAP), whose reduced levels are associated with neurodegeneration in amyotrophic lateral sclerosis and Parkinson’s disease (Kumar et al, 2018; Murphy and Levine, 2016; Slee and Levine, 2019).

### Properties of C-terminal VAB and PH-like domains

A C-terminal domain of VPS13 proteins was previously described as a tryptophan-aspartic acid (WD40) or a β-propeller-like domain depending on differently predicted structural folds (Dziurdzik et al, 2020; Kumar et al, 2018; Schapira et al, 2017). The VPS13 Adaptor Binding (VAB) domain lacks WD repeats in human VPS13s to be considered a WD40-like domain (PF00400), and in fact we did not find any significant structural similarities with a typical WD40 fold (Ye and Godzik, 2003). The VAB domain was previously described as a module containing six β-bladed repeats characterised by the presence of a highly conserved asparagine residue (**Video 2**; Bean et al, 2018; Dziurdzik et al, 2020; Dziurdzik and Conibear, 2021). Importantly, the VAB domain binds to organelle-specific interactors and was not expected to directly bind lipid membranes (Baldwin et al, 2021; Koike et al, 2019; Kumar et al, 2018; Muñoz-Braceras et al, 2019; Seifert et al, 2011; Yeshaw et al, 2019), as we show in our simulations (**Fig. 4; Videos S1-S4**). Interactions with proteins of the target organelle occur between the fifth or sixth VAB domain repeat and a consensus proline-x-proline motif (Bean et al, 2018) and likely contribute to the selectivity that VPS13C displays for endosomal rather than mitochondrial membrane binding *via* association with other proteins (Kumar et al, 2018).

At the C-terminus, an adenine phosphoribosyl transferase (APT1)-like domain was proposed to be present in yeast (Adlakha et al, 2022). However, we divide the previous APT1 and ATG-C domains into three subdomains in the VPS13 proteins. In all VPS13s except VPS13B (where the SGD is N-terminal to the VAP domain) the APT1 homology regions forms the SGD, while the β-sheets of the APT1 homology domain contribute to a narrowing hydrophobic β-sheet channel extending from the RBG repeats, terminated by a single amphipathic helix which is H1 of the membrane-bound gondola domain, shared with ATG2. This region is followed by a PH-like domain which generally interacts with phosphoinositides (Fidler et al, 2016). Extensively studied in yeast and human VPS13A, this PH-like domain was shown to bind Arf1 GTPase and *bis* and *tris*-phosphorylated phosphoinositides (Kolakowski et al, 2021). As our *in silico* approach and protein-membrane simulations did not predict the PH-like domains to exhibit significant hydrophobicity nor to be positively charged and to disturb lipid surfaces, suggesting membrane targeting interactions only (Videos S1-S4). The only exception is seen in VPS13C, where a loop extending from within the PH-like domain partially disturbs the surface of the endosomal membrane and may potentially compensate for the weak insertion of the VPS13C gondola (**Fig. 4**).

### Amphipathicity of VPS13 gondola domains

Amphipathic helices are membrane-targeting motifs that characterise the association of a variety of protein families with interactors and membranes (Drin and Antonny, 2010). Physicochemical properties such as hydrophobicity and the amount of net charge determine an amphipathic helix’s ability to target specific extrinsic domains or exhibit membrane lipid selectivity for binding (Prévost et al, 2018). From our VPS13 models and membrane simulations, it is possible to pinpoint the characteristics that allow the gondola domains to displace phospholipids. The gondola domain’s functional significance in lipid membrane binding is further emphasised by the presence of one hydrophobic and one hydrophilic wing (**Video 3**), one positively and one negatively electrostatic end (**Video 4**), and its highly conserved residues distributed homogenously over all surfaces of the gondola domains as we verified in conservation mapping (**Video 2**). Previous literature mentioned that the C-termini of VPS13 proteins share similar sequence identity to Dbl homology (DH) or RhoGEF domains consisting of ∼150-200 residues that generally allow GDP displacement and consist of an eleven α-helices bundle (Fig. 1; Kumar et al, 2018). However, the gondola domains contain 6-7 elongated helices which seem to be differently angled in pairs (**Fig 3**), much more similar to perilipins.

The mechanisms of lipid selectivity and cargo transfer regulation remain elusive. Nevertheless, we found that the gondola domains feature an α-helix pointing toward the interior of the RBG domain and this structural characteristic may direct cargoes reaching the C-terminal membrane leaflet, or alternatively, may block lipids enriched in the target membrane travelling ‘backwards’ to the donor membrane.

Homogenously positively charged strips on membrane-targeting protein structures are more likely to dive deeper within organelle membranes when these are rich in negatively charged lipids, as we established (**Videos S1-S4**). However, the negatively charged wing of the VPS13A gondola domain is also bound to the leaflet’s surface (Video S1, Video 4). It is possible that the presence of large hydrophobic residues sitting within the two helical bundles of the protein are the reason why the entire gondola domain, rather than only the positively charged wing, was predicted to associate with the leaflet. A fine balance between these properties is required to establish a functional anchor.

Particularly interesting is also that the hydrophobic and hydrophilic faces of the VPS13C gondola domain are inverted in orientation versus the membrane compared to those seen in VPS13B and VPS13D (**Video 3**). Only a small hydrophobic region of C-terminal VPS13C, indeed, binds the endosomal membrane, and this feature may, for instance, allow more space for a potential interactor to dock between the protein and the membrane. Thus, the VPS13C interactome and its strong binding affinity for Rab7 (McCray et al, 2010) may be what drives VPS13C’s preference for endolysosomal membranes (Kumar et al, 2018; Ramseyer et al, 2018; Yang et al, 2016) over those favoured by VPS13A.

Amphipathic helices are selective for certain membranes based on leaflet thickness, composition, and their conformation, as previously seen in lipid droplet surface-binding proteins (Prévost et al, 2018). Interestingly, these protein families display giant helices that may be up to 6 nm in length (**Fig. 3**). We discovered that the VPS13 gondola domains exhibit significant conformational similarities with the helical composition of ATG2A, Plin2, GPAT4 and, as such, contain similar extended-length helices curved at angles between 110-140° (Rowe et al, 2016).

Perilipins appear to associate at the membrane in a hierarchical fashion and hence the specificity and efficiency of lipid flux are regulated by the type of perilipins bound at the membrane (Ajjaji et al, 2019). The insertion of the amphipathic helices in perilipins serves several functions: one, to select membranes according to both headgroup and acyl chain composition, and the other allowing access to membrane lipids to other proteins by solubilizing these lipids, by bending and distorting the outer membrane leaflet (Sztalryd and Brasaemle, 2017). A similar mechanism may be at play in VPS13/ATG2 proteins: The same lipid flow and selectivity mechanisms could dictate cargo transfer in VPS13 proteins, as our modelling shows distinct structural features between VPS13A, VPS13B, VPS13C and VPS13D (**Fig. 4**). The gondola domain assumes very different geometries in the four human VPS13 proteins, suggesting selective association with membranes, and/or selective lipid transfer properties. However, in all four proteins, H1 of the gondola is aligned to the final β-sheet repeat of the RBG domain (Fig. 1), and extends the hydrophobic face of its amphipathic helix from the inner face of the pore to the surface of the membrane (Video 3). This may help conduct lipids to the outer leaflet of the target membrane, where the insertion of the gondola domain will distort the packing of lipids and most likely facilitate their movement and association with the organelle surface.

### Ideas and speculations

Overall, our modelling suggests that, although the Chorein domain inserts into donor membranes in our *in silico* models, the VPS13 N-terminus has a weaker association with the donor membrane compared to the VPS13 C-terminal domains where the gondola domain is inserted (**Fig. 4; Videos S1-S4**). *In vivo*, interactions at both ends of the protein are also facilitated and refined by interactions with resident proteins of each membrane. At the N-terminus this is mediated by an interaction with VAP proteins, which are anchored to the ER via a flexible peptide, and whose MSP domain contacts with the VPS13 FFAT motif, itself positioned in regions that are surrounded by unstructured residues at the seam of the RBG domain (Video 1). The C-terminal VAB and PH domains are known to interact with membrane lipids and small GTPases which help define target membrane identity, whereas the gondola mechanism is not particularly selective for membranes on its own (Kumar et al, 2018). The VPS13 C-terminus appears more geometrically constrained and in fact the PH-like and VAB domains here are already known to bind proteins labelling the target membrane (Adlakha et al, 2022; Kolakowski et al, 2021; McCray et al, 2010; Seifert et al, 2015). The role of the various SGDs and conserved peptide loops along the whole RBG length is as yet unknown despite similarities to known interaction motifs such as WWE and ricin B-type lectin domains (**Tables S1-S4**).

The crystallised Chorein domain provides a cavity that may accommodate several lipids at once, which is connected to a series of β-sheet repeats with a highly hydrophobic surface called the repeating β-groove (RBG) domain, which makes the bulk of the length of the protein (Fig. 1). These proteins appear to bind and transfer phosphatidylcholines and phosphatidylethanolamines (Kumar et al, 2018; Hanna et al, 2022) across intermembrane bridges formed by the RBG repeats, which is in line with our structure prediction models, where the highly hydrophobic inner channel of the RBG domains has an internal diameter of 4-6 nm that may accommodate several lipids at once (**Fig. 1**). This then narrows at the C terminus (1-2nm). To date, one key feature of all models of the RBG protein superfamily is that they adopt an open conformation where the highly hydrophobic inner surface is exposed to solvent (Neuman et al, 2022). Potentially, this may be necessary to facilitate or regulate lipid transport, or to confine lipid flow to a single ‘pathway’ through the RBG repeats. It is currently unknown how lipids are transported across RBG protein-formed bridges, if this transport is directional, and how this transport may be regulated. Recent studies show that directionality of transport is not necessarily an innate feature of all RBG domain-containing proteins, as SHIP164 forms a tail-to-tail dimer, where the Chorein domain is most likely embedded in both donor and target membranes (Hanna et al, 2022), and lipid transfer via SHIP164 appears to be between similar donor and target organelles. However, VPS13/ATG2 proteins span junctions between dissimilar organelles. Therefore, it is likely that specialised C terminal domains and the interactome of VPS13 proteins are necessary to impose a direction of transport which is not provided by the RBG repeats.

Recent evidence showed the RBG protein superfamily can transfer phosphatidylcholines and phosphatidylethanolamines (Neuman et al, 2022; Toulmay et al, 2022). The ER is relatively enriched in those lipid species compared to endo-lysosomes (Escribá et al, 2015; Guo et al, 2009; Horvath and Daum, 2013; Pogozheva et al, 2022; Reglinski et al, 2020; Tauchi-Sato et al, 2002; van Meer and de Kroon, 2011; van Meer et al, 2008). Despite this, the levels of phosphatidylcholine, phosphatidylserine, phosphatidylinositol, and sphingomyelin increase in the lysosome in VPS13C knockout clones (Hancock-Cerutti et al, 2022). Intriguingly, this suggests that VPS13C-mediated lipid transfer can be compensated for by other mechanisms.

*In vitro* experiments showed that the Chorein domain and several RBG repeats of yeast VPS13 are sufficient to extract phosphatidylethanolamine and/or phosphatidylserine from a ‘donor’ liposome and transfer this lipid to an acceptor liposome containing a lower concentration of both lipids (Kumar et al, 2018). This suggests passive diffusion, and no membrane-specific mechanism necessary for lipids to be accepted by the target *in vitro*. However, proteins like ATG2, which share the gondola domain with VPS13s, drive expansion of membrane compartments, which suggests asymmetric lipid transfer between donor and target. Our models highlight one possible mechanism that would establish donor-to-target directionality of flow. It is possible that ‘backwash’ i.e. the transfer of lipids enriched at the target membrane back to the donor membrane (here, the ER), might be prevented by the gondola embedded in the target membrane directly underneath the end of the narrowed C-terminal part of the hydrophobic channel conducting lipids.

Our modelling suggests that the C-terminus of VPS13s interacts more strongly with target membranes (in terms of depth of gondola penetration and hydrophobic surface of the C-terminus presented to the target membrane) than the N-terminus. This is despite *in vitro* experiments which dispense entirely with the VPS13C terminal domains to accomplish lipid transfer between model membranes. A stronger association of the VPS13 C terminus to target membranes compared to the N terminal Chorein domain to donor membranes is borne out by recent studies which measured the average density of the intramembrane bridges formed by VPS13C at contacts between the ER and lysosomes (Cai et al, 2022). In their recent paper, Cai et al produced a more tightly wound model which spanned 29.3 nm and left both ends of the protein free of membrane. However, their measurements in cells expressing full-length VPS13C showed the distribution of distances between ER and lysosome spanned by VPS13C centred around 32.5 nm (Cai et al, 2022), the exact length of VPS13C in our models (**Video 1**). Intriguingly, the density mapping of Cai et al (2022) showed that there was a substantial density on the C-terminal lysosomal side of the contact and little density where the protein was in contact via the chorein domain with the ER, suggesting a more stable conformation of the C terminus in their association with membranes than the N terminus.

This leads us to propose two possible mechanisms for VPS13-mediated lipid transfer, both anchored at the C-terminal target membrane, and assuming passive transfer of lipids down concentration gradients (**Fig. 5**). The first is a ‘piston’ model: the VPS13 proteins are relatively rigid down their length and interactors such as VAP proteins on the donor membrane, act to catch and draw VPS13 to contact with the organelle surface, allowing pulses of lipid transfer. The second is a ‘spring’ model, where the RBG repeats can adopt extended or more compressed geometry, changing the length of the interorganelle bridge. Conceivably, compression/relaxation of the spring could be facilitated by interactions with proteins of the intramembrane space (as yet unidentified) with the conserved cytosolic peptides and SGDs found along the RBG domain. Such a model sees the solvent exposure of the RBG interior changing as the spring compresses and relaxes, potentially regulating transfer of lipids with charged heads. In each model, the delivery of lipid to the target membrane would be driven by the insertion of the VPS13 gondola domain to its target, allowing addition of free lipids to the target membrane where the surface charge has been disturbed by the gondola to expose the hydrophobic chains of the outer leaflet (**Fig. 5**). This would then allow target membrane scramblases such as XK (Adlakha et al, 2022; Park and Neiman; 2020) to redistribute the transferred lipid to the correct leaflet of the target.

**Figure 5.**
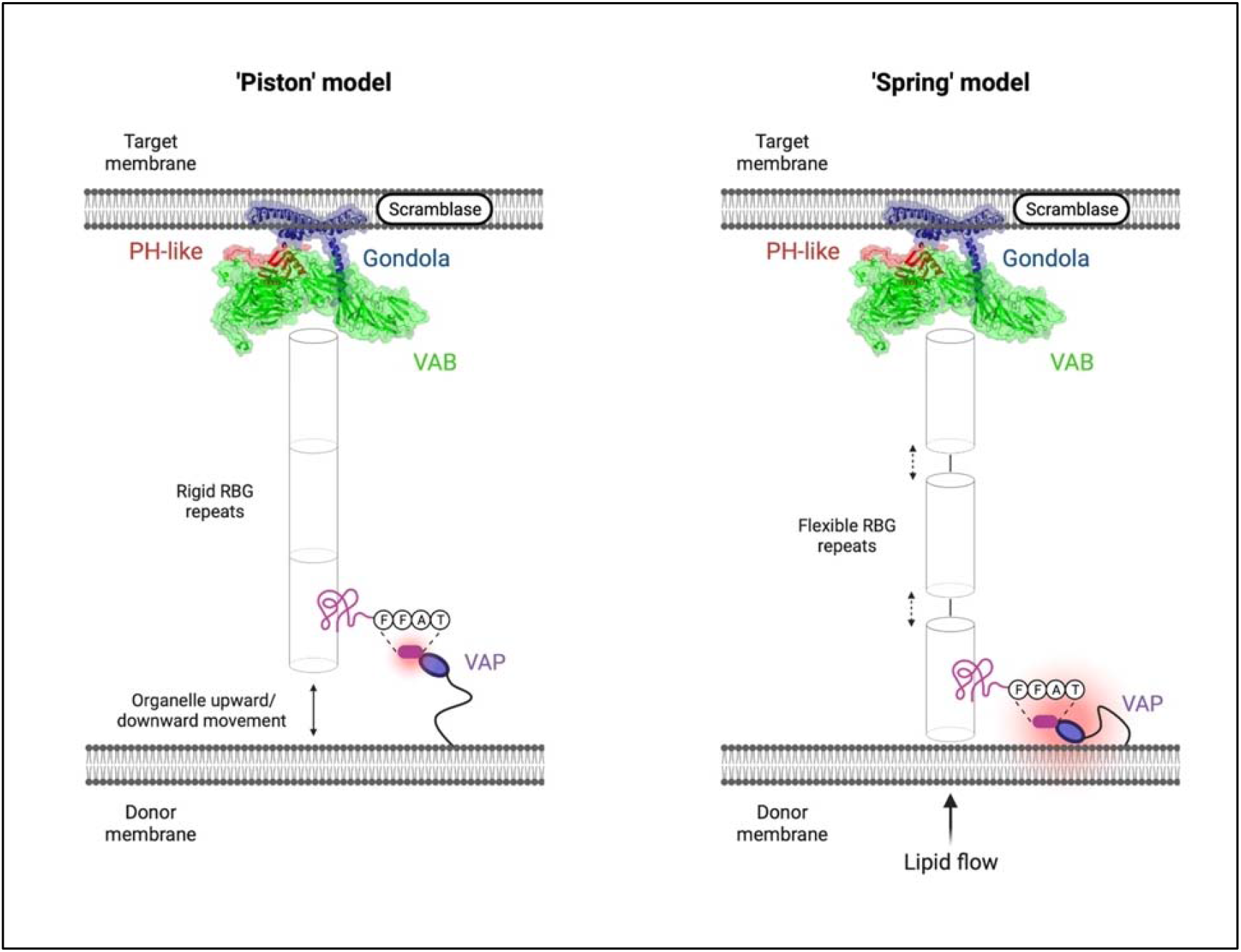
Speculative ‘piston’ and ‘spring’ models of VPS13-mediated intermembrane lipid transfer. The C-terminal end of the VPS13 protein in both models was taken from our modelling of VPS13D. The ‘piston’ model hypothesises that VPS13 proteins are rigidly docked at the target membrane *via* C-terminal interactions and the gondola domain and that the donor organelle performs upward/downward motions to free lipids for transfer. The ‘spring’ model hypothesises that lipid transfer occurs when both the N-terminus and C-terminus of VPS13 proteins are rigidly docked at their specific membranes and flow of transferring lipids is determined by compressing and relaxing the solvent-exposed gaps of the RBG domains. The figure was created on BioRender.com.

## Acknowledgements

We would like to acknowledge funding from Wellcome Trust ISSF 204822/Z/16/Z, MDUK to LES and AFM-Telethon to LES and MS.

---

**Video 1.**
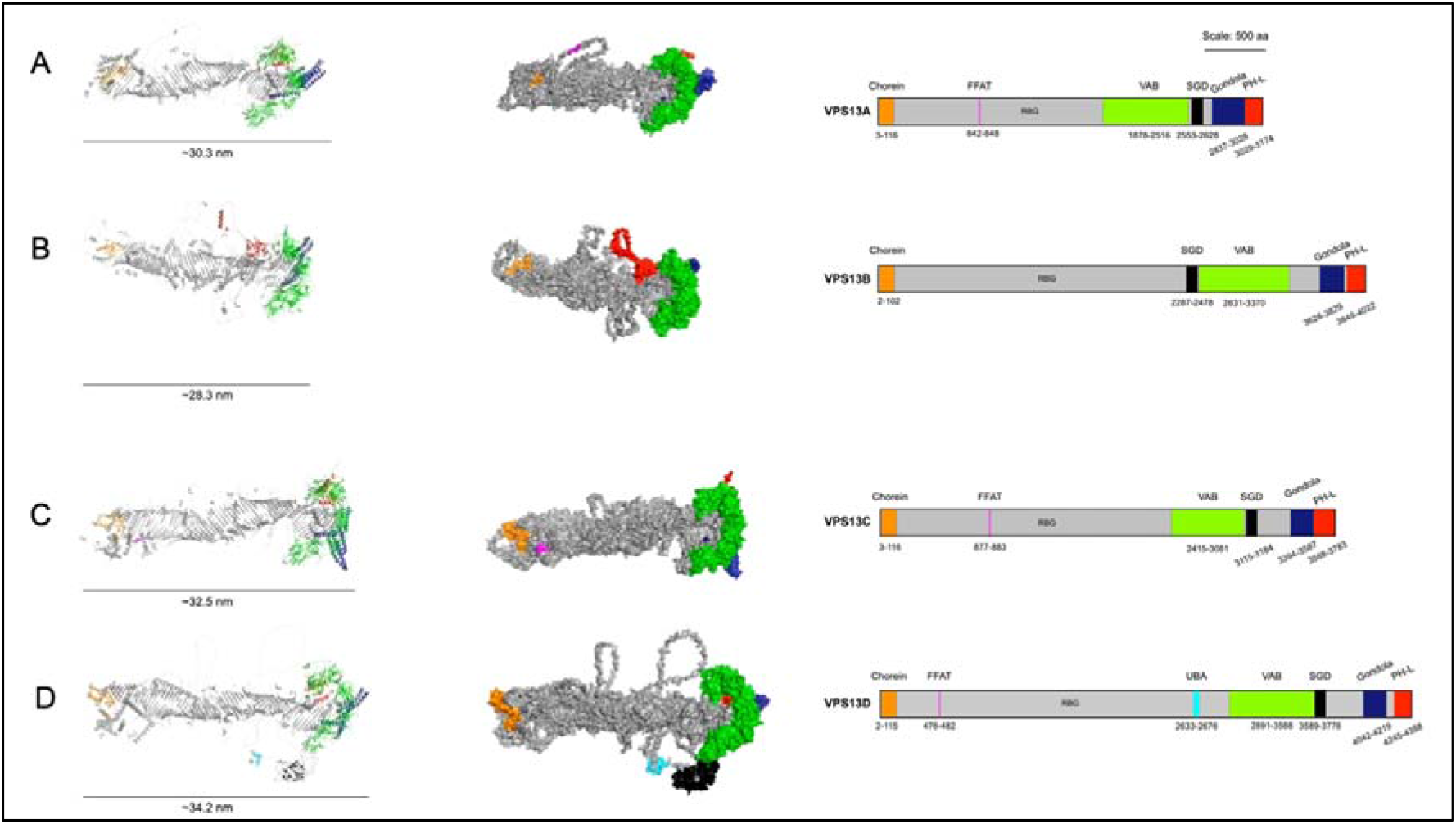
Full-atom structure prediction models of VPS13A, VPS13B, VPS13C, and VPS13D (A-D). Chorein domain (orange), FFAT motif (magenta), RBG domain (grey), UBA domain (azure), VAB domain (green), SGD (black), gondola domain (dark blue), PH-like domain (red). Model views were produced using PyMOL (Schrödinger and DeLano, 2020).

**Video 2.**
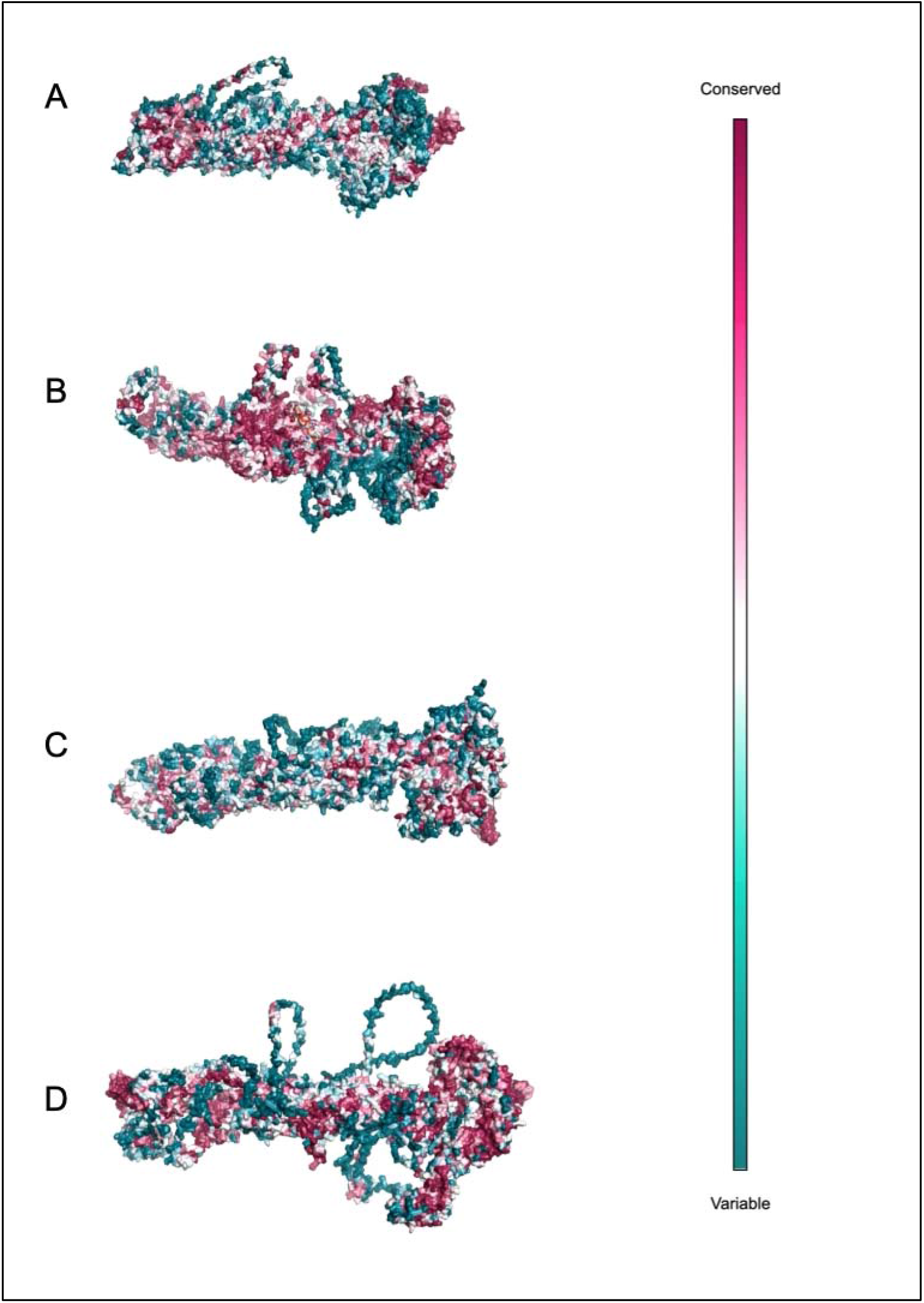
Conservation analysis of the human VPS13 proteins (A-D). The analysis was carried out employing the ConSurf Server (Ashkenazy et al, 2016; Landau et al, 2005). Darker purple-coloured residues are highly conserved and blue residues are variable in evolution.

**Video 3.**
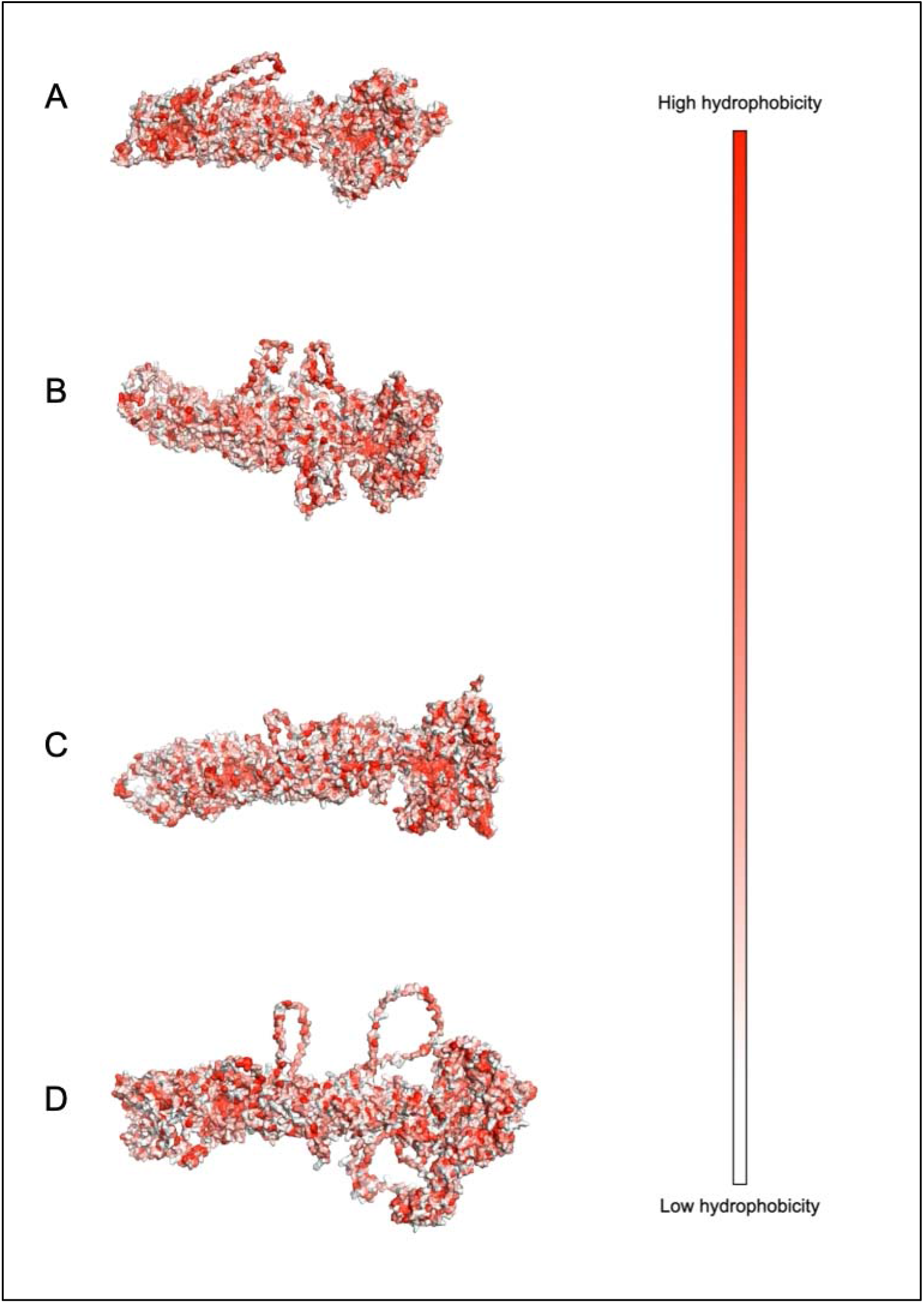
Hydrophobicity of the human VPS13 proteins (A-D). The Eisenberg scale (available at https://web.expasy.org/protscale/pscale/Hphob.Eisenberg.html), where highly hydrophobic residues are red and poorly hydrophobic are white.

**Video 4.**
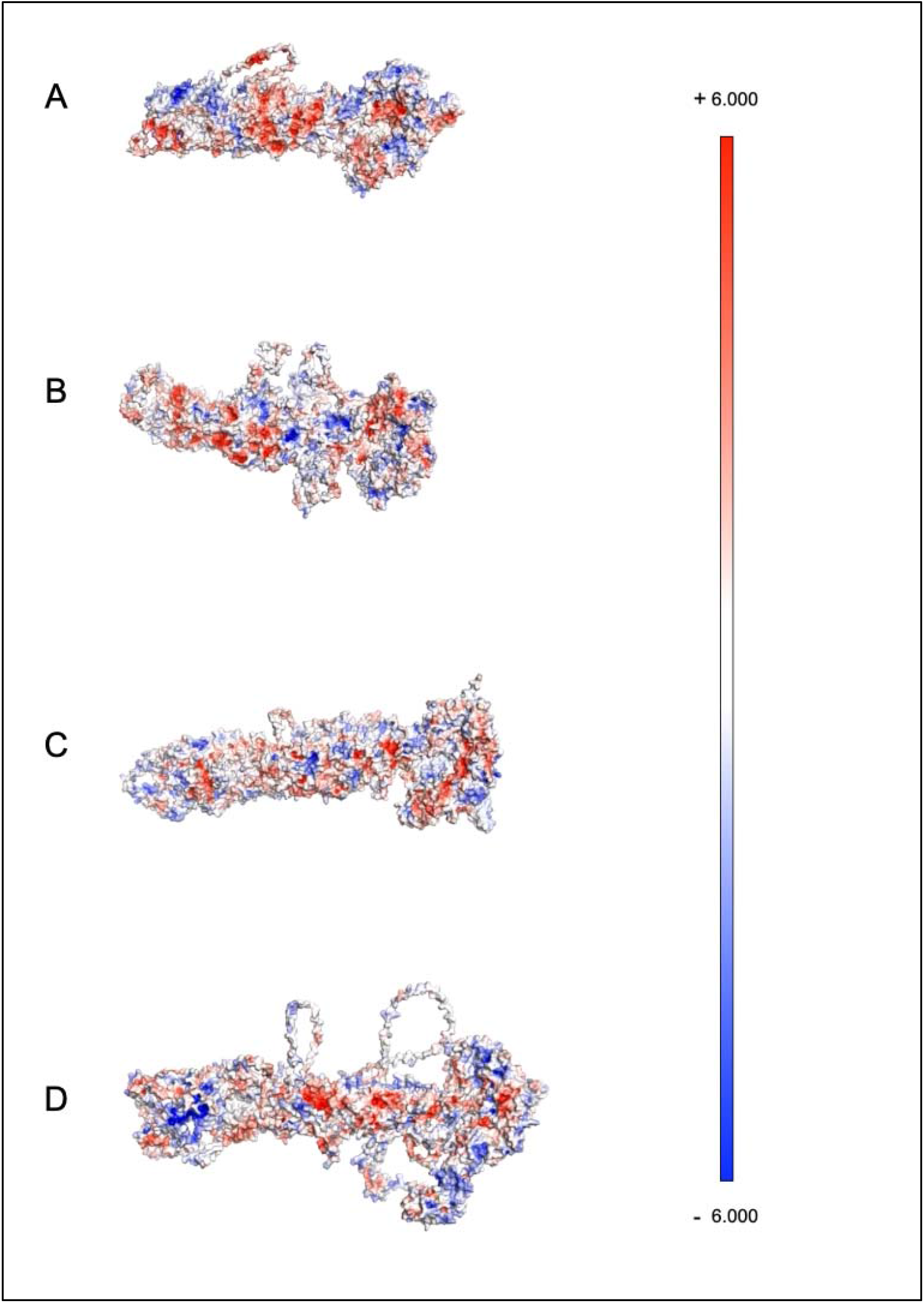
Electrostatic charge analysis of the human VPS13 proteins (A-D). Positively charged residues are red and negatively charged residues are blue, as calculated by the APBS Electrostatics Plugin in Incentive PyMOL 2.0 on the APBS-3.4.1 version of the software (Jurrus et al, 2018; Schrödinger and DeLano, 2020).

**Video S1.**
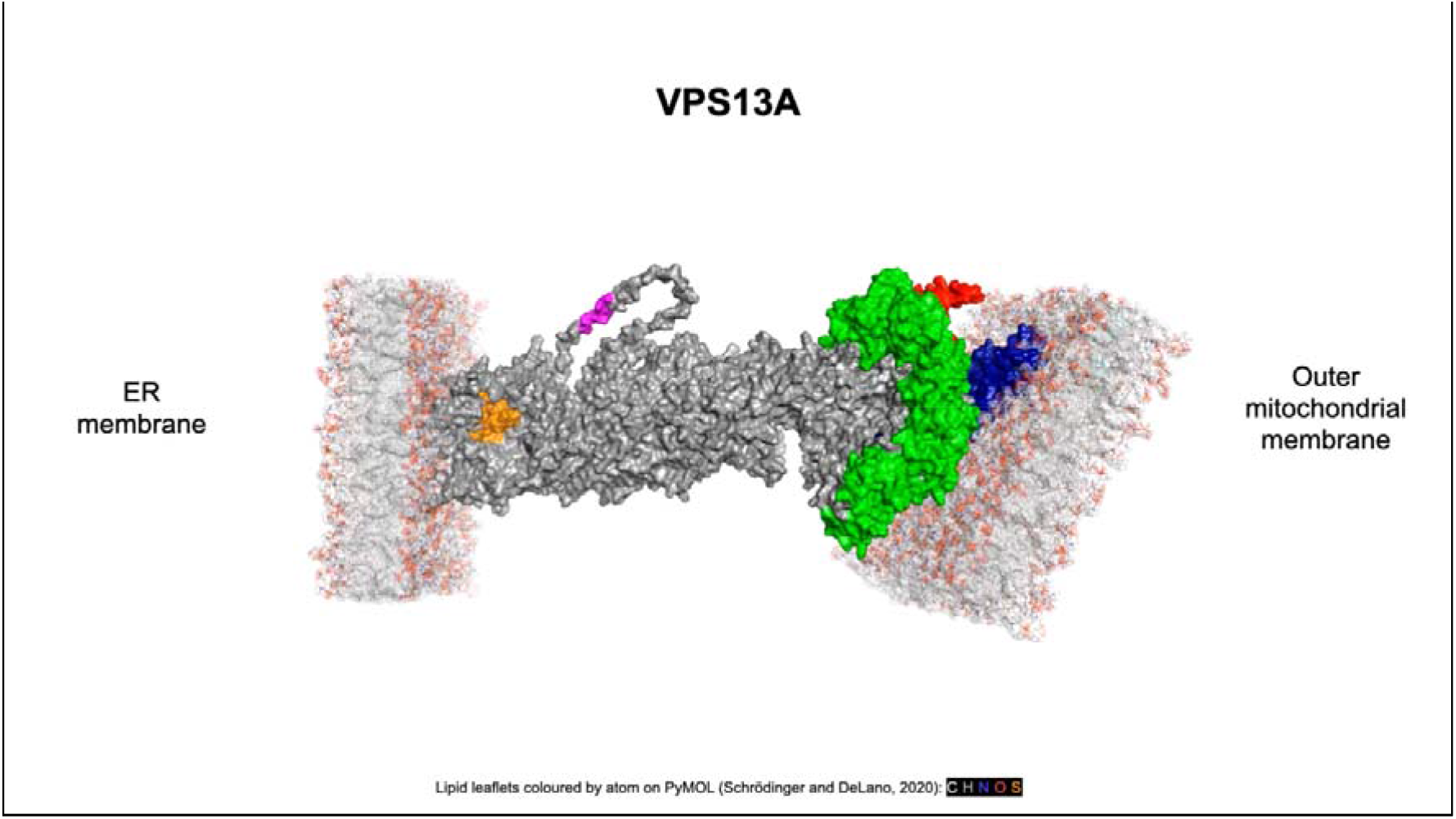
Simulation of human VPS13A at ER-mitochondria contact sites. *In silico* predictions were executed via CHARMM (Brooks et al, 2009; Jo et al, 2008, Wu et al, 2014). Chorein domain (orange), FFAT motif (magenta), RBG domain (grey), UBA domain (azure), VAB domain (green), SGD (black), gondola domain (dark blue), PH-like domain (red). Lipid leaflet compositions are reported in Tables S10-S18.

**Video S2.**
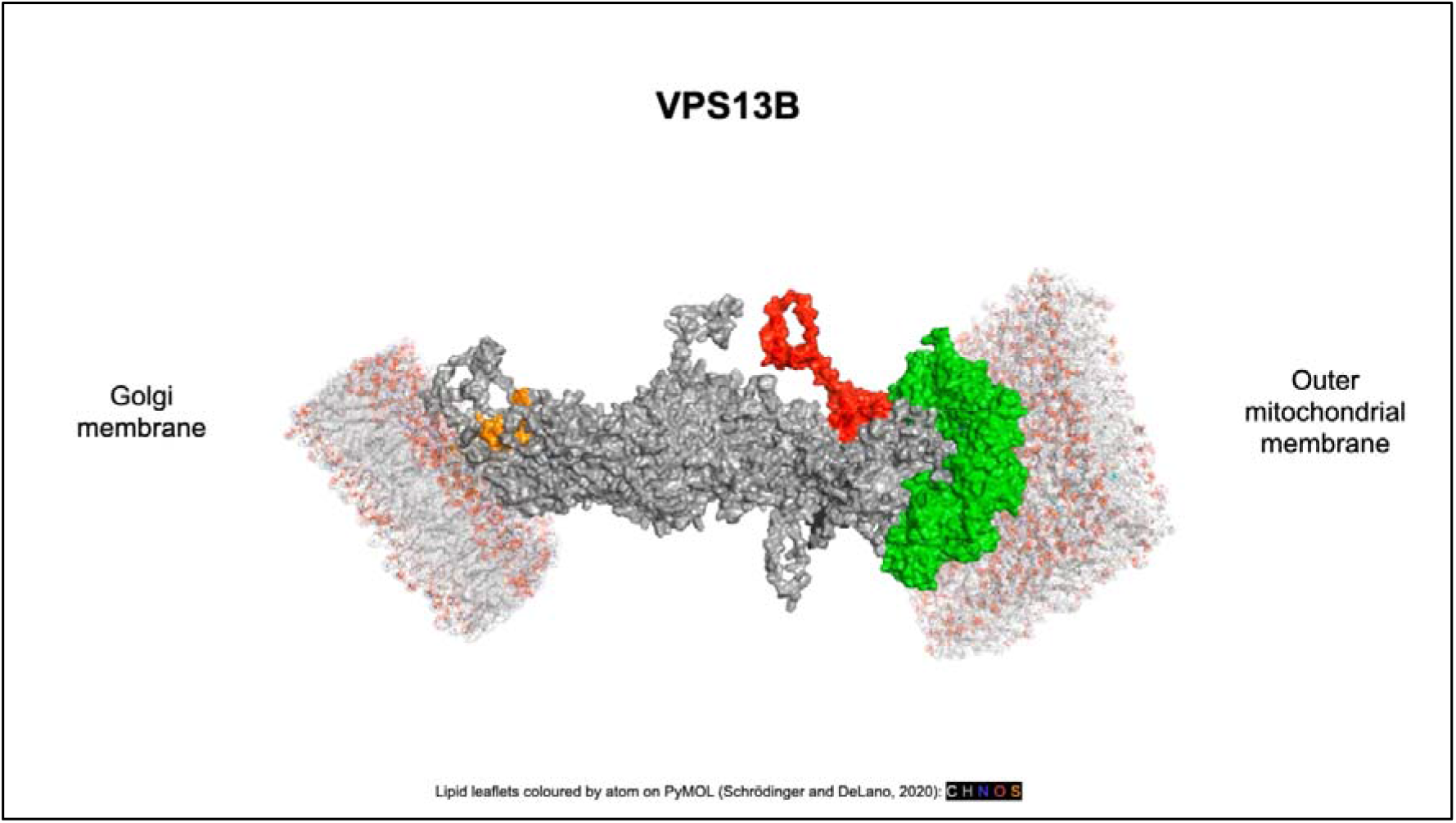
Simulation of human VPS13B at Golgi-mitochondria contact sites. *In silico* predictions were executed via CHARMM (Brooks et al, 2009; Jo et al, 2008, Wu et al, 2014). Chorein domain (orange), FFAT motif (magenta), RBG domain (grey), UBA domain (azure), VAB domain (green), SGD (black), gondola domain (dark blue), PH-like domain (red). Lipid leaflet compositions are reported in Tables S10-S18.

**Video S3.**
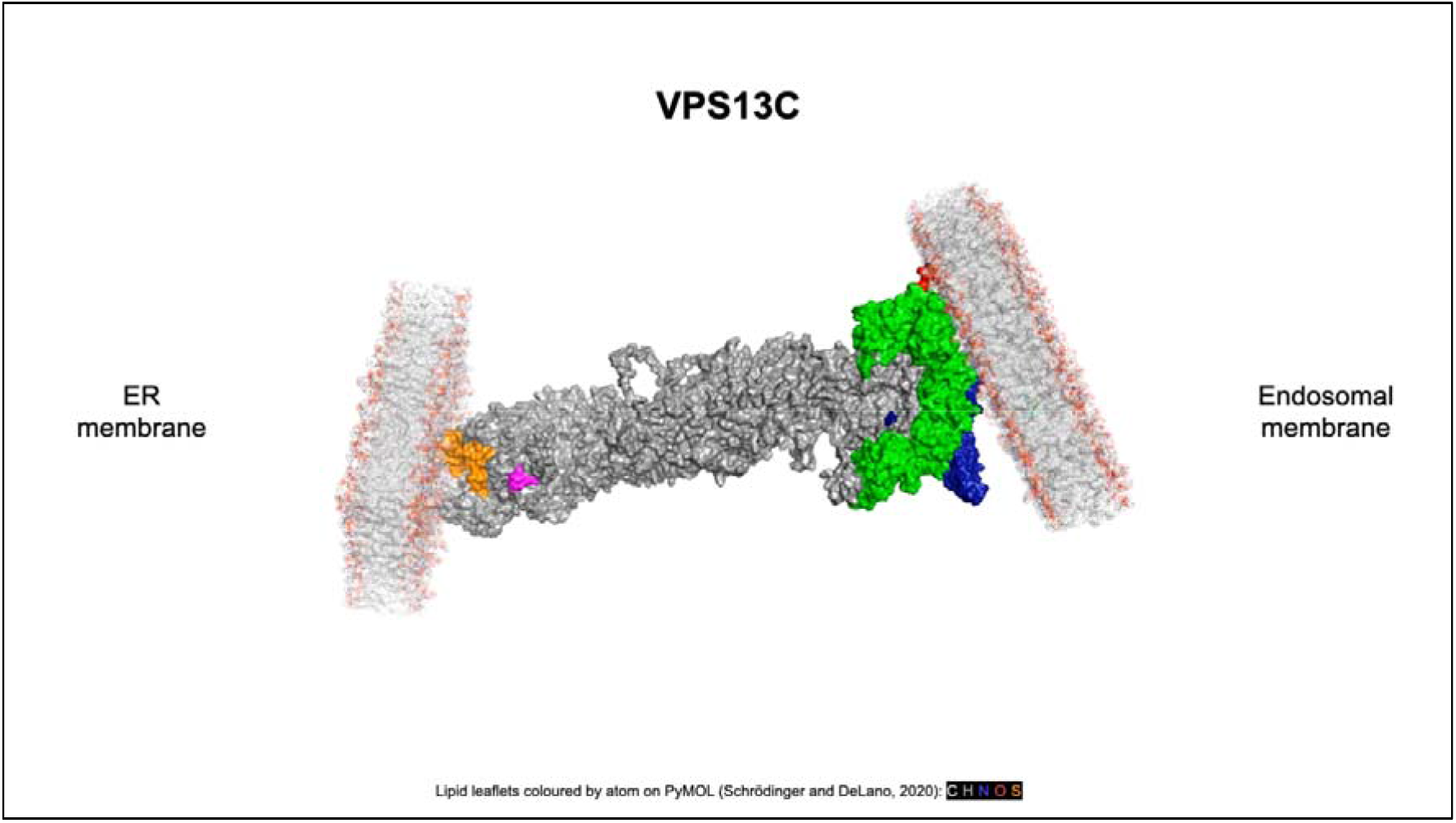
Simulation of human VPS13C at ER-endosome contact sites. *In silico* predictions were executed via CHARMM (Brooks et al, 2009; Jo et al, 2008, Wu et al, 2014). Chorein domain (orange), FFAT motif (magenta), RBG domain (grey), UBA domain (azure), VAB domain (green), SGD (black), gondola domain (dark blue), PH-like domain (red). Lipid leaflet compositions are reported in Tables S10-S18.

**Video S4.**
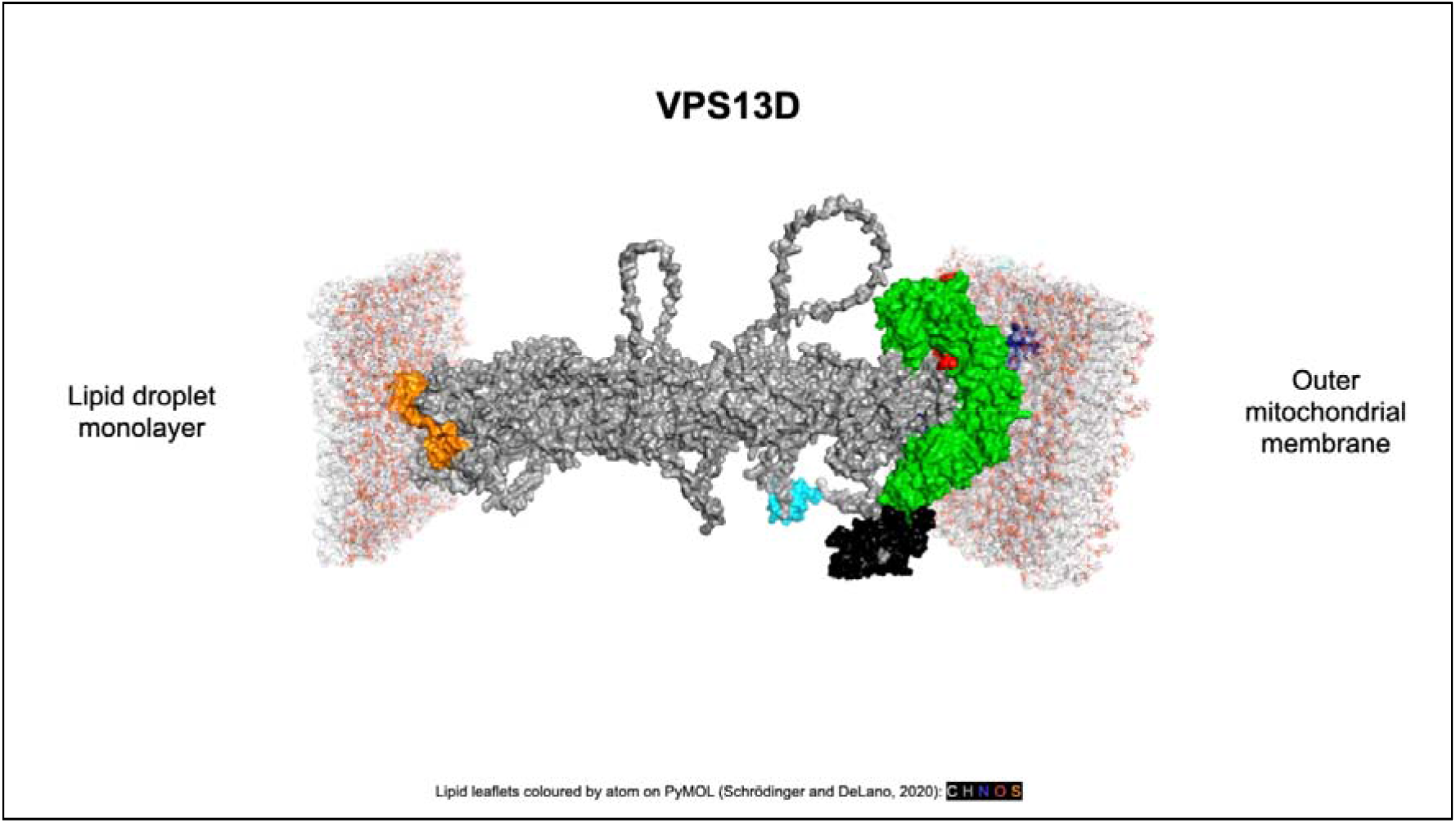
Simulation of human VPS13D at lipid droplet-mitochondria contact sites. *In silico* predictions were executed via CHARMM (Brooks et al, 2009; Jo et al, 2008, Wu et al, 2014). Chorein domain (orange), FFAT motif (magenta), RBG domain (grey), UBA domain (azure), VAB domain (green), SGD (black), gondola domain (dark blue), PH-like domain (red). Lipid leaflet compositions are reported in Tables S10-S18.

## Notes

### Competing Interest Statement

The authors have declared no competing interest.

