## supplementary material for "*In silico* modelling human VPS13 proteins associated with donor and target membranes suggests lipid transfer mechanisms"

Table S1. BackPhyre (curated) results for the submission of the VPS13A small globular domain against the Homo Sapiens genome database (Kelley et al, 2015).

| **Hit** | **Alignment coverage** | **Confidence** | **Percentage identity** |
| --- | --- | --- | --- |
| E3 ubiquitin-protein ligase DTX1 (NP_004407.2) | 1-68 | 98.3 | 20 |
| Deltex 2 (NP_065943.1) | 1-68 | 98.1 | 20 |
| SEC23-interacting protein (NP_009121.1) | 4-63 | 95.6 | 24 |
| DDHD domain containing 1 (NP_085140.1) | 4-50 | 92.7 | 17 |
| DDHD domain containing 2 (NP_056029.1) | 4-63 | 92.0 | 17 |

Table S2. BackPhyre (curated) results for the submission of the VPS13B small globular domain against the Homo Sapiens genome database (Kelley et al, 2015).

| **Hit** | **Alignment coverage** | **Confidence** | **Percentage identity** |
| --- | --- | --- | --- |
| N/A | N/A | N/A | N/A |

Table S3. BackPhyre (curated) results for the submission of the VPS13C small globular domain against the Homo Sapiens genome database (Kelley et al, 2015).

| **Hit** | **Alignment coverage** | **Confidence** | **Percentage identity** |
| --- | --- | --- | --- |
| Methanethiol oxidase (NP_003935.2) | 14-38 | 53.9 | 44 |

Table S4. BackPhyre (curated) results for the submission of the VPS13D small globular domain against the Homo Sapiens genome database (Kelley et al, 2015).

| **Hit** | **Alignment coverage** | **Confidence** | **Percentage identity** |
| --- | --- | --- | --- |
| GalNAc GALT-10 (NP_060010.3) | 28-126 | 97.7 | 27 |
| GalNAc-T12 (NP_078918.2) | 32-165 | 97.5 | 19 |
| Absent in melanoma 1-like protein (NP_001034864.1) | 18-126 | 97.5 | 20 |
| GALT-9 (NP_068580.2) | 32-144 | 97.5 | 22 |
| Beta/gamma crystallin domain-containing protein 3 (NP_705833.2) | 18-126 | 97.4 | 24 |
| GALT-17 (NP_001030017.1) | 28-127 | 97.2 | 27 |
| GALT-17 (NP_071924.1) | 32-156 | 97.1 | 20 |
| GALT-10 (NP_938080.1) | 28-126 | 97.0 | 26 |
| GALT-13 (NP_443149.1) | 23-127 | 96.9 | 31 |
| GALT-1 (NP_065207.2) | 23-126 | 96.8 | 39 |
| GALT-4 (NP_003765.2) | 32-178 | 96.4 | 14 |
| UDP-N-acetyl-alpha-D-galactosamine:GALT-2 (NP_473451.2) | 29-127 | 96.4 | 29 |
| GALT-2 (NP_004472.1) | 23-126 | 96.0 | 27` |
| GALT-14 (NP_078848.2) | 23-126 | 96.0 | 26 |
| GALNAC-T11 (NP_071370.1) | 32-165 | 95.2 | 21 |
| GALT-6 (NP_009141.1) | 32-143 | 95.2 | 12 |
| GALT-8 (NP_059113.1) | 26-126 | 94.9 | 19 |
| GALT-7 (NP_059119.1) | 23-126 | 94.8 | 19 |
| UDP-N-acetyl-alpha-D-galactosamine:GALT-4 (NP_940918.1) | 32-144 | 94.4 | 15 |
| GALT-5 (NP_055383.1) | 23-127 | 94.2 | 17 |
| GALT-3 (NP_004473.1) | 32-143 | 93.9 | 20 |
| PLA2R (NP_001007268.1) | 1-177 | 84.6 | 16 |
| GALT-1 (NP_065743.1) | 28-125 | 80.4 | 33 |
| PLA2R (NP_031392.3) | 1-177 | 78.7 | 16 |
| MRC1 (NP_002429.1) | 35-177 | 61.8 | 15 |

Table S5. BackPhyre (curated) results for the submission of the VPS13A gondola domain against the Homo Sapiens genome database (Kelley et al, 2015).

| **Hit** | **Alignment coverage** | **Confidence** | **Percentage identity** |
| --- | --- | --- | --- |
| ATG2A (NP_055919.1) | 2-191 | 100.0 | 18 |
| MTCH1 (NP_055156.1) | 28-170 | 75.1 | 24 |
| Plin-2 (NP_001113.2) | 24-188 | 62.3 | 11 |
| MPCPB (NP_998776.1) | 28-178 | 51.0 | 18 |

Table S6. BackPhyre (curated) results for the submission of the VPS13B gondola domain against the Homo Sapiens genome database (Kelley et al, 2015).

| **Hit** | **Alignment coverage** | **Confidence** | **Percentage identity** |
| --- | --- | --- | --- |
| ATG2A (NP_055919.1) | 2-200 | 100.0 | 19 |
| MTCH1 (NP_055156.1) | 28-179 | 56.8 | 15 |

Table S7. BackPhyre (curated) results for the submission of the VPS13C gondola domain against the Homo Sapiens genome database (Kelley et al, 2015).

| **Hit** | **Alignment coverage** | **Confidence** | **Percentage identity** |
| --- | --- | --- | --- |
| ATG2A (NP_055919.1) | 2-193 | 100.0 | 19 |

Table S8. BackPhyre (curated) results for the submission of the VPS13D gondola domain against the Homo Sapiens genome database (Kelley et al, 2015).

| **Hit** | **Alignment coverage** | **Confidence** | **Percentage identity** |
| --- | --- | --- | --- |
| ATG2A (NP_055919.1) | 2-174 | 99.9 | 18 |
| MTCH1 (NP_055156.1) | 29-153 | 73.2 | 16 |
| MTCH2 (NP_055157.1) | 29-169 | 64.8 | 13 |

Table S9. Distribution and functional significance of pathogenic missense mutations in human VPS13 proteins (National Center for Biotechnology Information, ClinVar; https://www.ncbi.nlm.nih.gov/clinvar, accessed July 24, 2022). The nomenclature may vary from the ClinVar database reports as it was adapted to the canonical transcripts used for modelling.

| **VPS13A (NP_150648.2)** | | | |
| --- | --- | --- | --- |
| **Nomenclature** | **Location** | **Topological orientation** | 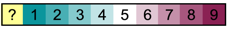**Conservation (0-9)** |
| [p.Ile90Lys](https://www.ncbi.nlm.nih.gov/clinvar/variation/4682/) | Chorein β-sheet | Groove inward-facing | 6 |
| [p.Ala2679Thr](https://www.ncbi.nlm.nih.gov/clinvar/variation/4686/) | β-sheet between SGD-Gondola | Cytosolic-facing | 6 |
| **VPS13B (NP_060360.3)** | | | |
| [p.Met1Ile](https://www.ncbi.nlm.nih.gov/clinvar/variation/402220/) | Pre-Chorein loop | Cytosolic-facing | 9 |
| [p.Gln27Arg](https://www.ncbi.nlm.nih.gov/clinvar/variation/813329/) | Chorein β-sheet | Cytosolic-facing | 9 |
| [p.Pro1133Ser](https://www.ncbi.nlm.nih.gov/clinvar/variation/225021/) | RBG loop | Cytosolic-facing | 6 |
| [p.Ile1636Asn](https://www.ncbi.nlm.nih.gov/clinvar/variation/977843/) | RBG β-sheet | Inward-facing | 9 |
| [p.Ser1812Ile](https://www.ncbi.nlm.nih.gov/clinvar/variation/427100/) | RBG loop | Cytosolic-facing | 6 |
| [p.Leu2193Arg](https://www.ncbi.nlm.nih.gov/clinvar/variation/2819/) | RBG β-sheet | Inward-facing | 4 |
| [p.Leu2543Val](https://www.ncbi.nlm.nih.gov/clinvar/variation/975775/) | RBG β-sheet | Cytosolic/SGD-facing | 2 |
| **VPS13C (NP_065872.1)** | | | |
| [p.Gly1389Arg](https://www.ncbi.nlm.nih.gov/clinvar/variation/222070/) | RBG loop | Cytosolic-facing | 1 |
| **VPS13D (NP_056193.2)** | | | |
| [p.Ile1170Ser](https://www.ncbi.nlm.nih.gov/clinvar/variation/806071/) | RBG β-sheet | Inward-facing | 9 |
| [p.Gly1190Asp](https://www.ncbi.nlm.nih.gov/clinvar/variation/561198/) | RBG loop | Cytosolic-facing | 6 |
| [p.Asn3521Ser](https://www.ncbi.nlm.nih.gov/clinvar/variation/561203/) | VAB domain β-sheet | Cytosolic-facing | 6 |
| [p.Ala4139Val](https://www.ncbi.nlm.nih.gov/clinvar/variation/932894/) | Gondola domain H4 | Membrane-bound | 8 |
| [p.Gly4177Asp](https://www.ncbi.nlm.nih.gov/clinvar/variation/983263/) | Gondola domain hinge between H5-H6 | Membrane-bound | 9 |
| [p.Ala4210Val](https://www.ncbi.nlm.nih.gov/clinvar/variation/561200/) | Gondola domain H6 | Membrane-bound | 9 |
| [p.Arg4228Gln](https://www.ncbi.nlm.nih.gov/clinvar/variation/561204/) | Loop between gondola and PH-like domains | Cytosolic-facing | 6 |
| [p.Ala4248Glu](https://www.ncbi.nlm.nih.gov/clinvar/variation/807716/) | PH-like domain α-helix | Cytosolic/VAB-facing | 9 |

Table S10. Protein-membrane system calculations between the N-terminus (1-215 aa) of VPS13A and an ER membrane (Brooks et al, 2009; Jo et al, 2008, Wu et al, 2014).

| **Calculated Lipid Number** | | |
| --- | --- | --- |
| **Lipid type** | **Upper-leaflet number** | **Lower-leaflet number** |
| Cholesterol | 15 | 15 |
| POPA | 3 | 3 |
| POPC | 48 | 48 |
| PLPC | 54 | 54 |
| PSPE | 24 | 24 |
| POPE | 15 | 15 |
| OLPS | 9 | 9 |
| SLPI | 9 | 9 |
| SAPI | 9 | 9 |
| SAPC | 81 | 81 |
| SAPE | 21 | 21 |
| PSM | 12 | 12 |
| **Calculated XY system size** | | |
|  | **Upper-leaflet** | **Lower-leaflet** |
| Protein Area | 0 | 657.992985 |
| Lipid area | 19369.8 | 19369.8 |
| Number of Lipids | 300 | 300 |
| Total area | 19369.8 | 20027.792985 |
| Protein X extent | 33.42 | |
| Protein Y extent | 31.86 | |
| Average area | 19698.80 | |
| A | 140.35 | |
| B | 140.35 | |

Table S11. Protein-membrane system calculations between the C-terminus (1876-3174 aa) of VPS13A and an outer mitochondrial membrane (Brooks et al, 2009; Jo et al, 2008, Wu et al, 2014).

| **Calculated Lipid Number** | | |
| --- | --- | --- |
| **Lipid type** | **Upper-leaflet number** | **Lower-leaflet number** |
| POPA | 3 | 3 |
| POPC | 18 | 18 |
| PLPC | 54 | 54 |
| POPE | 6 | 12 |
| PLPE | 15 | 27 |
| SAPI | 66 | 12 |
| TLCL1 | 0 | 6 |
| SAPC | 90 | 90 |
| SAPE | 39 | 75 |
| SAPS | 9 | 3 |
| **Calculated XY system size** | | |
|  | **Upper-leaflet** | **Lower-leaflet** |
| Protein Area | 0 | 1798.08463 |
| Lipid area | 20153.1 | 20317.5 |
| Number of Lipids | 300 | 300 |
| Total area | 20153.1 | 22115.58463 |
| Protein X extent | 71.44 | |
| Protein Y extent | 58.24 | |
| Average area | 21134.34 | |
| A | 145.38 | |
| B | 145.38 | |

Table S12. Protein-membrane system calculations between the N-terminus (1-205 aa) of VPS13B and a Golgi complex membrane (Brooks et al, 2009; Jo et al, 2008, Wu et al, 2014).

| **Calculated Lipid Number** | | |
| --- | --- | --- |
| **Lipid type** | **Upper-leaflet number** | **Lower-leaflet number** |
| Cholesterol | 24 | 24 |
| POPC | 33 | 33 |
| PLPC | 42 | 42 |
| PSPE | 24 | 24 |
| PLPE | 12 | 12 |
| POPS | 12 | 12 |
| PSPI | 6 | 6 |
| POPI | 15 | 15 |
| SAPI | 6 | 6 |
| SAPC | 60 | 60 |
| SAPE | 15 | 15 |
| TSM | 36 | 36 |
| LPC16 | 15 | 15 |
| **Calculated XY system size** | | |
|  | **Upper-leaflet** | **Lower-leaflet** |
| Protein Area | 0 | 989.62037 |
| Lipid area | 18298.2 | 18298.2 |
| Number of Lipids | 300 | 300 |
| Total area | 18298.2 | 19287.82037 |
| Protein X extent | 45.07 | |
| Protein Y extent | 22.33 | |
| Average area | 18793.01 | |
| A | 137.09 | |
| B | 137.09 | |

Table S13. Protein-membrane system calculations between the C-terminus (2630-4022 aa) of VPS13B and an outer mitochondrial membrane (Brooks et al, 2009; Jo et al, 2008, Wu et al, 2014).

| **Calculated Lipid Number** | | |
| --- | --- | --- |
| **Lipid type** | **Upper-leaflet number** | **Lower-leaflet number** |
| POPA | 3 | 3 |
| POPC | 18 | 18 |
| PLPC | 54 | 54 |
| POPE | 6 | 12 |
| PLPE | 15 | 27 |
| SAPI | 66 | 12 |
| TLCL1 | 0 | 6 |
| SAPC | 90 | 90 |
| SAPE | 39 | 75 |
| SAPS | 9 | 3 |
| **Calculated XY system size** | | |
|  | **Upper-leaflet** | **Lower-leaflet** |
| Protein Area | 0 | 2319.90896 |
| Lipid area | 20153.1 | 20317.5 |
| Number of Lipids | 300 | 300 |
| Total area | 20153.1 | 22637.40896 |
| Protein X extent | 63.27 | |
| Protein Y extent | 84.56 | |
| Average area | 21395.25 | |
| A | 146.27 | |
| B | 146.27 | |

Table S14. Protein-membrane system calculations between the N-terminus (1-225 aa) of VPS13C and an ER membrane (Brooks et al, 2009; Jo et al, 2008, Wu et al, 2014).

| **Calculated Lipid Number** | | |
| --- | --- | --- |
| **Lipid type** | **Upper-leaflet number** | **Lower-leaflet number** |
| Cholesterol | 15 | 15 |
| POPA | 3 | 3 |
| POPC | 48 | 48 |
| PLPC | 54 | 54 |
| PSPE | 24 | 24 |
| POPE | 15 | 15 |
| OLPS | 9 | 9 |
| SLPI | 9 | 9 |
| SAPI | 9 | 9 |
| SAPC | 81 | 81 |
| SAPE | 21 | 21 |
| PSM | 12 | 12 |
| **Calculated XY system size** | | |
|  | **Upper-leaflet** | **Lower-leaflet** |
| Protein Area | 0 | 630.23516 |
| Lipid area | 19369.8 | 19369.8 |
| Number of Lipids | 300 | 300 |
| Total area | 19369.8 | 20000.03516 |
| Protein X extent | 52.74 | |
| Protein Y extent | 24.90 | |
| Average area | 19684.92 | |
| A | 140.30 | |
| B | 140.30 | |

Table S15. Protein-membrane system calculations between the C-terminus (2444-3783 aa) of VPS13C and an endosomal membrane (Brooks et al, 2009; Jo et al, 2008, Wu et al, 2014).

| **Calculated Lipid Number** | | |
| --- | --- | --- |
| **Lipid type** | **Upper-leaflet number** | **Lower-leaflet number** |
| PLPC | 10 | 10 |
| SAPC | 16 | 16 |
| SDPC | 6 | 6 |
| PLPE | 2 | 2 |
| PDoPE | 4 | 4 |
| SAPE | 6 | 6 |
| SAPI | 6 | 6 |
| SDPI | 2 | 2 |
| DSM | 8 | 8 |
| OSM | 8 | 8 |
| Cholesterol | 32 | 32 |
| BMGP | 280 | 280 |
| **Calculated XY system size** | | |
|  | **Upper-leaflet** | **Lower-leaflet** |
| Protein Area | 0 | 489.54831 |
| Lipid area | 23849 | 23849 |
| Number of Lipids | 380 | 380 |
| Total area | 23849 | 24338.54831 |
| Protein X extent | 68.14 | |
| Protein Y extent | 54.98 | |
| Average area | 24093.77 | |
| A | 155.22 | |
| B | 155.22 | |

Table S16. Protein-membrane system calculations between the C-terminus (2444-3783 aa) of VPS13C and an outer mitochondrial membrane (Brooks et al, 2009; Jo et al, 2008, Wu et al, 2014).

| **Calculated Lipid Number** | | |
| --- | --- | --- |
| **Lipid type** | **Upper-leaflet number** | **Lower-leaflet number** |
| POPA | 4 | 4 |
| POPC | 24 | 24 |
| PLPC | 72 | 72 |
| POPE | 8 | 16 |
| PLPE | 20 | 36 |
| SAPI | 88 | 16 |
| TLCL1 | 0 | 8 |
| SAPC | 120 | 120 |
| SAPE | 52 | 100 |
| SAPS | 12 | 4 |
| **Calculated XY system size** | | |
|  | **Upper-leaflet** | **Lower-leaflet** |
| Protein Area | 0 | 489.54831 |
| Lipid area | 23984.8 | 23984.8 |
| Number of Lipids | 382 | 382 |
| Total area | 23984.8 | 24474.34831 |
| Protein X extent | 68.14 | |
| Protein Y extent | 54.98 | |
| Average area | 24229.57 | |
| A | 155.66 | |
| B | 155.66 | |

Table S17. Protein-membrane system calculations between the N-terminus (1-225 aa) of VPS13D and a lipid droplet monolayer (Brooks et al, 2009; Jo et al, 2008, Wu et al, 2014).

| **Calculated Lipid Number** | | |
| --- | --- | --- |
| **Lipid type** | **Upper-leaflet number** | **Lower-leaflet number** |
| Cholesterol | 15 | 15 |
| POPA | 3 | 3 |
| POPC | 48 | 48 |
| PLPC | 54 | 54 |
| PSPE | 24 | 24 |
| POPE | 15 | 15 |
| OLPS | 9 | 9 |
| SLPI | 9 | 9 |
| SAPI | 9 | 9 |
| SAPC | 81 | 81 |
| SAPE | 21 | 21 |
| PSM | 12 | 12 |
| **Calculated XY system size** | | |
|  | **Upper-leaflet** | **Lower-leaflet** |
| Protein Area | 0 | 1049.68063 |
| Lipid area | 19369.8 | 19369.8 |
| Number of Lipids | 300 | 300 |
| Total area | 19369.8 | 20419.48063 |
| Protein X extent | 33.48 | |
| Protein Y extent | 30.34 | |
| Average area | 19894.64 | |
| A | 141.05 | |
| B | 141.05 | |

Table S18. Protein-membrane system calculations between the C-terminus (2889-4388 aa) of VPS13D and an outer mitochondrial membrane (Brooks et al, 2009; Jo et al, 2008, Wu et al, 2014).

| **Calculated Lipid Number** | | |
| --- | --- | --- |
| **Lipid type** | **Upper-leaflet number** | **Lower-leaflet number** |
| POPA | 3 | 3 |
| POPC | 18 | 18 |
| PLPC | 54 | 54 |
| POPE | 6 | 12 |
| PLPE | 15 | 27 |
| SAPI | 66 | 12 |
| TLCL1 | 0 | 6 |
| SAPC | 90 | 90 |
| SAPE | 39 | 75 |
| SAPS | 9 | 3 |
| **Calculated XY system size** | | |
|  | **Upper-leaflet** | **Lower-leaflet** |
| Protein Area | 0 | 1673.81361 |
| Lipid area | 20153.1 | 20317.5 |
| Number of Lipids | 300 | 300 |
| Total area | 20153.1 | 21991.31361 |
| Protein X extent | 97.00 | |
| Protein Y extent | 57.57 | |
| Average area | 21072.21 | |
| A | 145.16 | |
| B | 145.16 | |
